# Breakage of the Oligomeric CaMKII Hub by the Regulatory Segment of the Kinase

**DOI:** 10.1101/2020.04.15.043067

**Authors:** Deepti Karandur, Moitrayee Bhattacharyya, Beryl Xia, Young Kwang Lee, Serena Muratcioglu, Darren McAffee, Ethan D. McSpadden, Baiyu Qiu, Jay Groves, Evan Williams, John Kuriyan

## Abstract

Ca^2+^/calmodulin dependent protein kinase II (CaMKII) is a dodecameric or tetradecameric enzyme with crucial roles in neuronal signaling and cardiac function. Activation of CaMKII is reported to trigger the exchange of subunits between holoenzymes, which can increase spread of the active state. Using mass spectrometry, we now show that peptides derived from the sequence of the CaMKII-α regulatory segment can bind to the CaMKII-α hub assembly and break it into smaller oligomers. Molecular dynamics simulations show that the regulatory segments can dock spontaneously at the interface between hub subunits, trapping large fluctuations in hub structure. Single-molecule fluorescence intensity analysis of human CaMKII-α isolated from mammalian cells shows that activation of CaMKII-α results in the destabilization of the holoenzyme. Our results show how the release of the regulatory segment by activation and phosphorylation could allow it to destabilize the hub, producing smaller CaMKII assemblies that can reassemble to form new holoenzymes.

## Introduction

Ca^2+^/calmodulin-dependent protein kinase II (CaMKII) is an oligomeric serine/threonine kinase that is important in neuronal signaling and cardiac function (Bhattacharyya et al., 2019; Kennedy, 2013; Lisman et al., 2002). Each subunit of CaMKII has an N-terminal kinase domain that is followed by a regulatory segment and an unstructured linker that leads into a C-terminal hub domain (Figure 1A, B). The regulatory segment blocks the substrate-binding site of the kinase in the autoinhibited state of CaMKII (Rellos et al., 2010; Rosenberg et al., 2005), and this inhibition is released by the binding of Ca^2+^/calmodulin (Ca^2+^/CaM) to a calmodulin-binding element within the regulatory segment (Figure 1B) (Ikura et al., 1992; Meador et al., 1993, 1992; Rellos et al., 2010). The hub domains of CaMKII associate to form a ring-shaped scaffold containing twelve or fourteen subunits, around which the kinase domains are arranged (Figure 1A) (Chao et al., 2011; Hoelz et al., 2003; Myers et al., 2017; Rellos et al., 2010). Mutations in CaMKII affect learning and memory formation in mice (Elgersma et al., 2002; Giese et al., 1998; Silva et al., 1992), and mutations in human CaMKII have been identified in patients with cognitive and developmental impediments (Chia et al., 2018; Küry et al., 2017; Robison, 2014).

**Figure 1.**
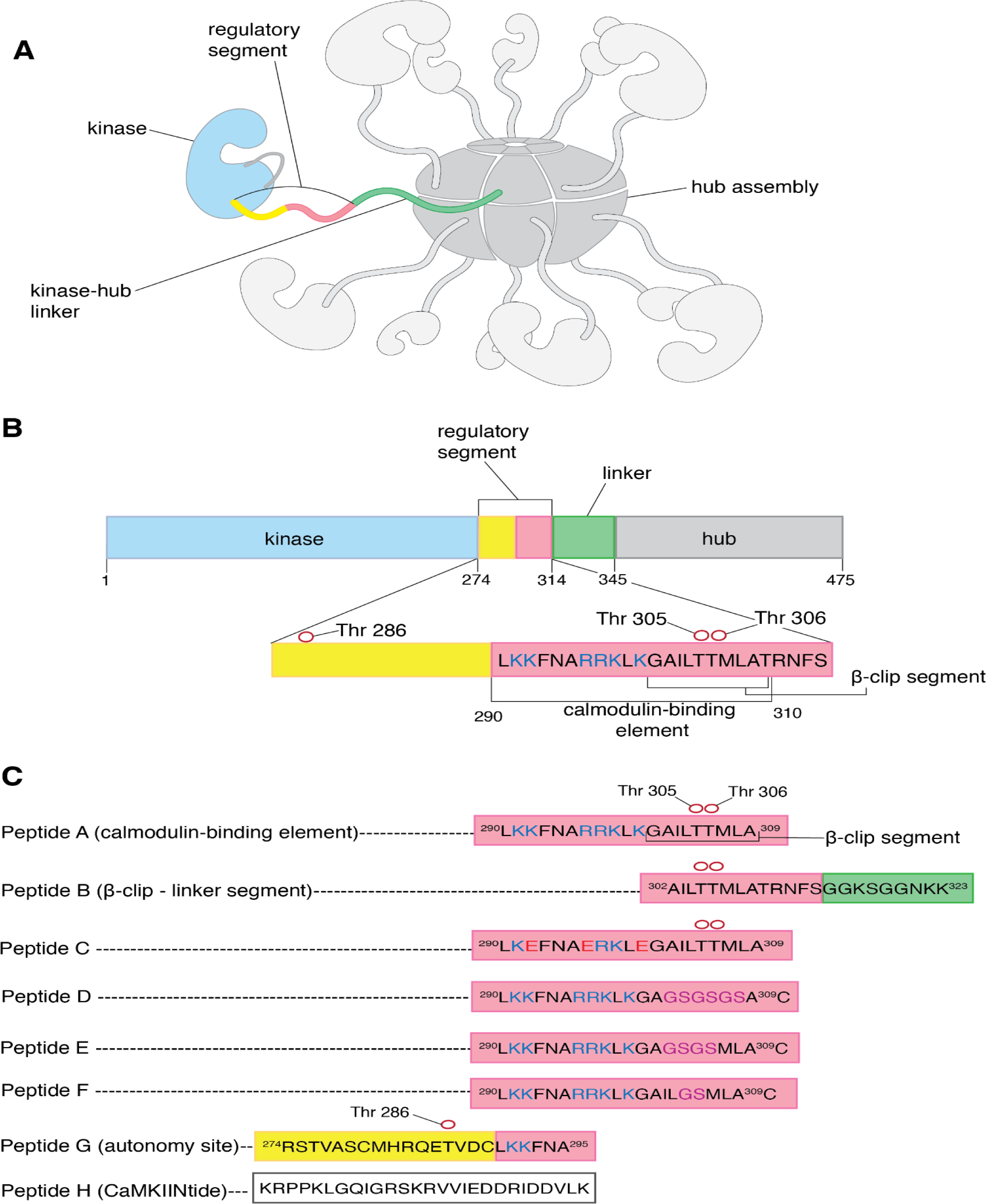
CaMKII architecture and peptides used in mass spectrometry analysis. (**A)** The architecture of dodecameric CaMKII, with twelve subunits. The kinase domains are followed by the regulatory segments, which are connected to the hub domains by linkers. The hub domains (in darker grey) associate to form a ring-like hub assembly. For clarity, only a single kinase domain, regulatory segment, and linker are shown in colors, while the remainder are shown in shades of grey. In this subunit, the kinase domain is colored pale blue, the regulatory segment is colored yellow and pale pink, and the unstructured kinase-hub linker is colored green. **(B)** Domains of a CaMKII-α subunit. The color scheme is the same as in **A**. The calmodulin-binding element and the β-clip segment are labeled. The positively charged residues of the calmodulin-binding element are colored deep blue. Phosphorylation sites are shown as empty red circles. **(C)** Peptides used in the mass spectrometry analyses. All the peptides are derived from the sequence of CaMKII-α, except Peptide H, a peptide that inhibits the kinase domain of CaMKII. The color schemes denote the region of the CaMKII-α subunit that the peptide is derived from and are the same as in **A**. All the peptides, except Peptide H, are aligned with respect to Peptide A. Positively charged residues of the calmodulin-binding element are colored blue, and the β-clip segment is labeled in Peptide A. Phosphorylation sites in Peptide A and Peptide G are shown as empty red circles. In Peptide C positively charged residues that are replaced with negatively charged residues are colored red. In Peptides D, E and F residues that are replaced with glycine and serine residues are colored purple.

The activation of CaMKII by Ca^2+^/calmodulin triggers the co-localization of subunits from different CaMKII holoenzymes, indicating that activation results in subunits being exchanged between holoenzymes (Bhattacharyya et al., 2016; Stratton et al., 2014). This conclusion was based on the results of experiments in which two samples of CaMKII were labeled separately with fluorophores of two different colors, mixed, and then activated. Single-molecule visualization showed activation-dependent co-localization of the two fluorophores (Stratton et al., 2014). In other experiments, separate CaMKII samples were labeled with FRET donor and acceptor fluorophore pairs, respectively, and mixing these samples after activation led to increased FRET compared to mixing unactivated samples (Bhattacharyya et al., 2016). These data are also consistent with subunit exchange. Colocalization of differently-labeled subunits was also observed when unactivated CaMKII holoenzymes were mixed with activated ones. Importantly, these experiments showed that activated CaMKII holoenzymes could phosphorylate subunits of unactivated ones, thereby spreading the activation signal (Stratton et al., 2014).

The regulatory segment of CaMKII has been shown to be important for subunit exchange (Bhattacharyya et al., 2016; Stratton et al., 2014). After activation, autophosphorylation of Thr 286 in the regulatory segment prevents it from re-binding to the kinase domain, thereby conferring Ca^2+^/CaM independence (autonomy) on CaMKII (Lou and Schulman, 1989; Miller et al., 1988; Thiel et al., 1988). Autophosphorylation of two other sites, Thr 305 and Thr 306, prevents the binding of Ca^2+^/CaM to the regulatory segment (Colbran, 1993). Thus, we expect that activation is accompanied by the release of at least some fraction of the regulatory segment from the kinase and from Ca^2+^/CaM.

A constitutively-active variant of CaMKII-α (T286D) undergoes spontaneous subunit exchange without activation by Ca^2+^/CaM (Stratton et al., 2014). Mutation of the calmodulin-binding element of the regulatory segment in this variant eliminated subunit exchange, which suggests that the calmodulin-binding element might interact with the hub and destabilize it, thereby promoting subunit exchange (Stratton et al., 2014). Consistent with this, a construct consisting of just the hub and the regulatory segment was shown to undergo spontaneous subunit exchange (Bhattacharyya et al., 2016).

A prominent feature of the hub assembly is the presence of grooves located at the inter-subunit interfaces. These grooves are lined by negatively-charged residues, and we proposed that positively-charged residues in the calmodulin-binding element might recognize these negatively-charged residues (Bhattacharyya et al., 2016). Each interfacial groove between subunits contains the uncapped edge of a β-sheet, and peptide segments have been shown to dock within the groove by forming an additional strand of this β-sheet (Bhattacharyya et al., 2016; Chao et al., 2011). A peptide with the sequence of the calmodulin-binding element has been shown to bind weakly to the hub, with a dissociation constant of ∼90 µM, but destabilization of the hub was not studied in those experiments (Bhattacharyya et al., 2016).

In this paper, we present the results of investigations into interactions between the regulatory segment and the hub, and the effect of activation on the stability of the holoenzyme. We used electrospray ionization mass spectrometry to show that peptides derived from the regulatory segment can bind to and destabilize the dodecameric or tetradecameric CaMKII-α hub assembly, releasing smaller oligomers of the hub domain. We generated a 13 μs molecular dynamics trajectory of the dodecameric hub of CaMKII-α, which shows that two regulatory segments bind spontaneously to the hub at two inter-subunit interfaces. This docking is facilitated by large distortions in the structure of the hub, which reflect an intrinsic feature of hub dynamics, as shown by a normal mode calculation. Finally, we use single-molecule TIRF microscopy to show that activation of CaMKII-α expressed in mammalian cells is accompanied by apparent destabilization of the holoenzyme.

## Results and Discussion

### Mass spectrometry shows that peptides derived from the regulatory segment can break the CaMKII-α hub assembly

We used electrospray ionization mass spectrometry to analyze the integrity of the CaMKII-α hub assembly, in isolation and in the presence of peptides. The sequences of most of these peptides are based on that of the calmodulin-binding element of the regulatory segment (Peptide A and Peptides C-F, Figure 1B), which includes several positively-charged residues preceding hydrophobic residues. We refer to the hydrophobic portion (residues 301-308) as the β-clip segment because a part of this segment is incorporated into the interfacial β-sheet in a structure of autoinhibited CaMKII-α, in a conformation referred to as the β-clip conformation (Chao et al., 2011). We also tested a peptide that contains only the β-clip segment and a portion of the kinase-hub linker (Peptide B). Finally, we tested the effect of two different peptides that bind to the kinase domain of CaMKII but have no known affinity for the hub. One of these peptides (Peptide G) corresponds to the portion of the regulatory segment that precedes the calmodulin-binding element (residues 274-295). This region of the regulatory segment binds to the kinase domain in the autoinhibited state and includes Thr 286, which confers Ca^2+^/CaM-independent activity, or “autonomy” when phosphorylated (Lou and Schulman, 1989). The other peptide (Peptide H) is a commonly used CaMKII inhibitor called CaMKIINtide, which binds to the kinase domain and blocks the active site (Chang et al., 1998; Chao et al., 2010).

The mass spectrometry experiments were carried out for the CaMKII-α hub in isolation, without the kinase domain and linkers. Intact CaMKII-α holoenzymes yield poor quality mass spectra due to sample heterogeneity resulting from proteolysis during the purification process (Bhattacharyya et al., 2016). The hub (120 µM subunit concentration) was incubated, with or without peptides, for five minutes at room temperature prior to injection into a time-of-flight mass spectrometer under soft ionization conditions. The resulting ions were detected, and the masses of the different species present in the sample were determined (see Methods for complete description). The intact hub is detected in the *m*/*z* range of 5800 to 8000, and smaller oligomeric species appear in the *m*/*z* range of 2000 to 5200.

The mass spectrum of the CaMKII-α hub without added peptides is shown in Figure 2A. We observe a distribution of ions only in the higher *m*/*z* range, which correspond to dodecameric and tetradecameric hub assemblies. These observations are similar to the results of previous native mass spectrometry studies on isolated CaMKII-α hubs, in which dodecameric and tetradecameric hub assemblies were observed (Bhattacharyya et al., 2016; McSpadden et al., 2019). In the previous studies, a higher sample cone voltage of 150 V was used to reduce adduction of solvent and/or other small species, in order to obtain higher-resolution mass spectra. Under these higher-energy source conditions, some collision-induced disassembly of the hub occurs, resulting in the release of a small amount of monomeric species (Figure 2SA). In the present study, we used softer ionization conditions, with a cone voltage of 50 V. Although this results in a decrease in the mass spectral resolution, we do not observe any smaller oligomeric species for the isolated hub. These acquisition conditions are therefore preferable for distinguishing between stable hub assemblies and those with peptide-induced disassembly, and were used for the remainder of the experiments.

**Figure 2.**
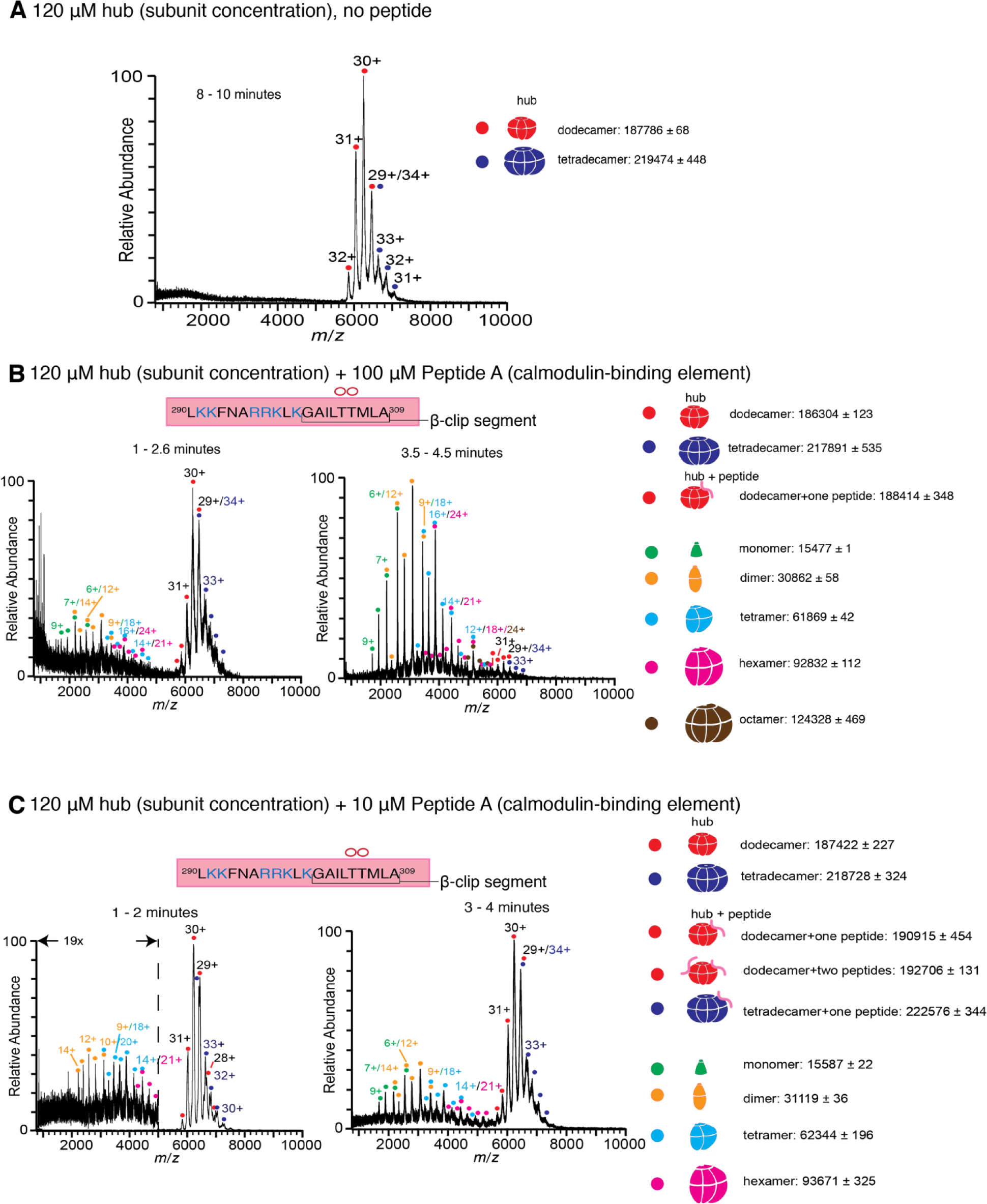
Peptides derived from the calmodulin-binding element of CaMKII-α bind to the hub assembly and result in its disassembly. **(A)** The isolated hub assembly is comprised of dodecamers and tetradecamers. **(B)** The hub assembly (120 µM subunit concentration) incubated with 100 µM of Peptide A, shows disassembly within 3 minutes of ionization (left). In addition to dodecamers and tetradecamers, monomers, dimers, tetramers, hexamers and octamers are observed. At ∼4 minutes (right), the smaller oligomers are predominant. **(C)** The hub assembly (120 µM subunit concentration) incubated with 10 µM of Peptide A shows some release of smaller oligomeric species at ∼2 minutes (left). This becomes more pronounced at 3-4 minutes (right), although the dodecameric and tetradecameric hub assemblies are also present. In all the spectra of the hub incubated with peptides, species corresponding to dodecameric or tetradecameric hub assemblies with one or two peptides bound are present.

Peptide A, containing the calmodulin-binding element, was added at final concentrations of 1 µM, 10 µM, 100 µM and 1 mM to CaMKII-α hub at a final subunit concentration of 120 µM. Following incubation, the sample was ionized and mass spectra were generated by averaging the scans for specific time segments after the initiation of electrospray ionization. The dissociation constant of Peptide A for the hub is ∼90 µM (Bhattacharyya et al., 2016), and so we expect appreciable binding of the peptide to the hub for the condition with 100 µM of Peptide A. Indeed, the mass spectrometry results with 100 µM of Peptide A and the hub are dramatically different than for the hub alone (Figure 2B). In the presence of 100 µM of Peptide A, we observe hub species comprised of dodecamers and tetradecamers, as well as new species corresponding in mass to the intact complex with a peptide adducted. In addition, smaller oligomeric species that correspond to monomers, dimers, tetramers and hexamers of hub subunits appear within less than 3 minutes (Figure 2B). After 4 minutes these smaller species are predominant.

Incubation of the hub with Peptide A at 10 µM concentration also yields spectra that demonstrate the release of smaller oligomeric species within ∼1-2 minutes, although the peptide is sub-stoichiometric with respect to the hub under this mixing condition (Figure 2C). At ∼3-4 minutes the smaller species are prominent, although the dodecameric species are also present (Figure 2C). At even lower concentrations (1 µM) of Peptide A, we see essentially no disassembly of the hub, even after ∼10 minutes, although there is some hexamer present marginally above the noise. The abundance of peptide-adducted complex is significantly reduced at this lower peptide concentration. We note that under these conditions, the ratio of peptide to hub subunits is ∼1:100 (Figure 2SB). Incubation at saturating concentrations of peptide (1 mM) yields very noisy spectra with a high baseline consistent with significant protein/peptide aggregation (Figure 2SC).

The incubation of the hub with 100 µM of Peptide B also causes disassembly of the hub and the release of monomers, dimers, tetramers and hexamers, but at a slower rate than seen with Peptide A. There are no smaller species observed up to 2.5 minutes (Figure 3A); smaller oligomeric species are detectable only at ∼3-4 minutes (Figure 3A). Peptide B does not contain the positively-charged residues of the calmodulin-binding element. To test the importance of positive charge in Peptide A, we replaced three positively-charged residues with glutamate (Peptide C, see Figure 1B). Addition of 1 mM of Peptide C to the hub did not cause hub disassembly (Figure 3B). These results are in agreement with previously reported binding measurements that showed that this peptide fails to bind to the hub (Bhattacharyya et al., 2016).

**Figure 3.**
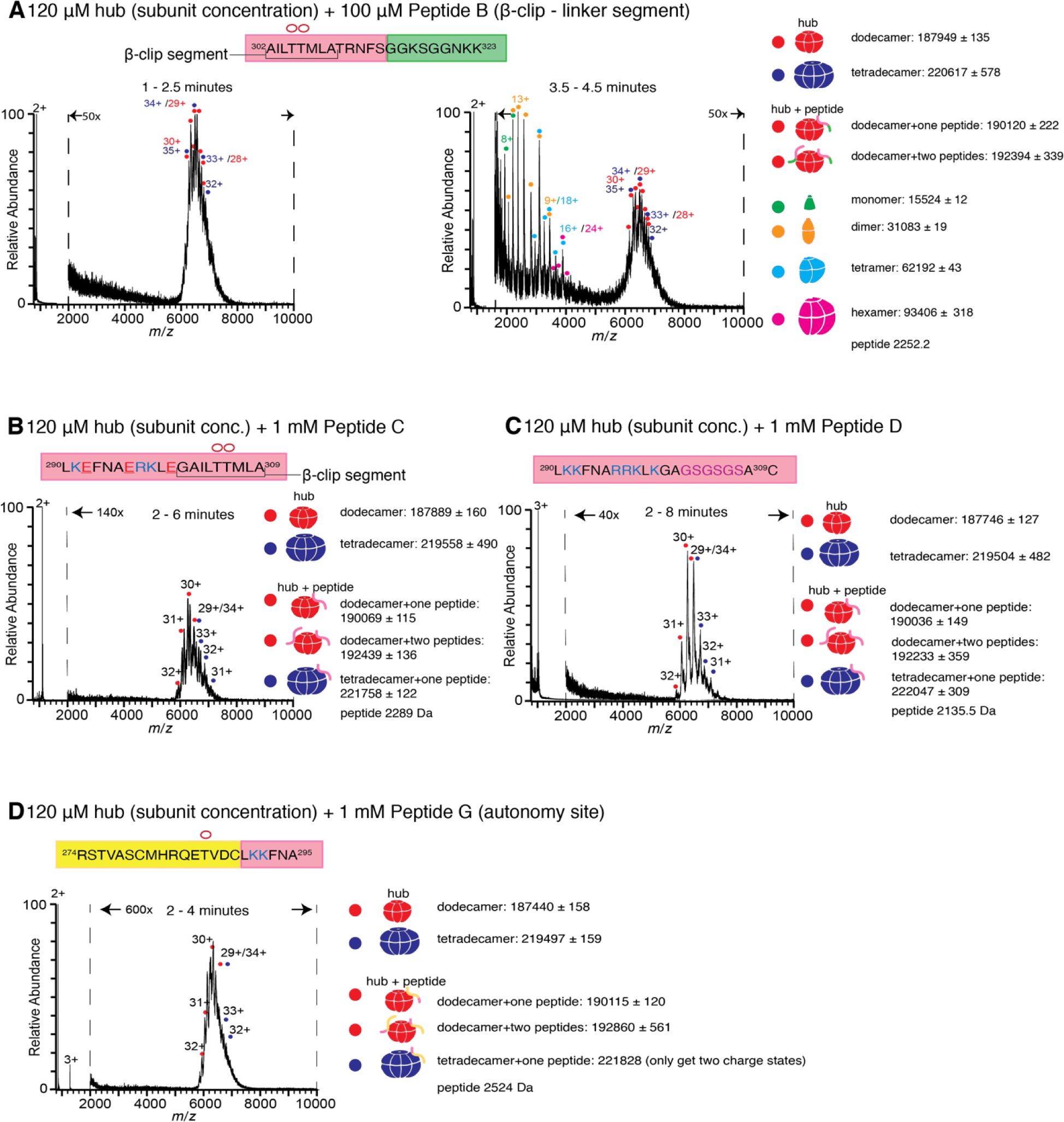
Different regions of the calmodulin-binding element of CaMKII-α play roles in inducing hub disassembly. **(A)** Peptide B, which contains the β-clip segment of the calmodulin-binding element, but not the positively charged residues induces disassembly, but at a slower rate than Peptide A. When 100 µM of Peptide B is incubated with the hub assembly, no smaller species are observed up to ∼2.5 minutes after ionization (left). At 3.5-4.5 minutes, smaller oligomeric species corresponding to monomers, dimers, tetramers and hexamers are observed (right). **(B)** Incubation of the hub assembly with 1 mM of Peptide C, in which three positively-charged residues of Peptide A are replaced with negatively-charged residues (colored red in sequence) does not result in the release of smaller oligomeric species up to 6 minutes after ionization, even though the peptide binds to the hub assembly. **(C)** Modifying the β-clip segment of the calmodulin-binding element abrogates disassembly. Incubation of the hub assembly with 1 mM of Peptide D, in which the β-clip segment of Peptide A is replaced with glycine and serine residues (colored purple) does not result in the release of smaller oligomeric species, and only dodecamers and tetradecamers are observed up to 8 minutes after ionization. **(D)** Incubation of the hub assembly with 1 mM of Peptide G, whose sequence corresponds to the autonomy site of the CaMKII-α regulatory segment, does not result in the release of smaller oligomeric species up to 4 minutes after ionization.

We replaced the β-clip segment of Peptide A with glycine and serine residues (Peptides D, E and F, see Figure 1B). Modification of the β-clip segment reduces or eliminates the ability of the peptides to disturb the integrity of the hub. The results of the incubation with 1 mM of Peptide D, where the entire β-clip segment is replaced with glycine and serine residues, are shown in Figure 3C. Only intact dodecamers and tetradecamers are observed after 8 minutes. Peptides E and F, in which only portions of the β-clip segment are replaced by glycine and serine residues, cause hub disassembly (Figure 3SA, 3SB), but to a lesser extent than Peptide A at similar peptide concentrations (Figure 2SC).

We also tested the effect of two control peptides that are known to interact with the kinase domain, but are not known to bind to the hub. The first (Peptide G, Figure 1B) contains the autonomy site (Thr 286) of the regulatory segment (Lou and Schulman, 1989), and the second, CaMKIINtide (Peptide H, Figure 1B), is an inhibitor that binds to the substrate recognition site of the kinase domain (Chang et al., 1998; Chao et al., 2010). Some adduction of these peptides to the intact complexes is observed, but neither peptide induces hub disassembly (Figure 3D, 3SC), pointing to a specific role for the calmodulin-binding element in disturbing the integrity of the CaMKII-α hub.

### Long-timescale molecular dynamics simulations of regulatory segment docking on the hub

We generated molecular dynamics trajectories for the hub assembly with the regulatory segments and linkers either present or absent, using the specialized Anton2 supercomputer (Shaw et al., 2014). We built a dodecameric hub assembly of CaMKII-α, based on the crystal structure of the CaMKII-α hub (PDB ID – 5IG3) (Bhattacharyya et al., 2016), on to which we modeled the unstructured linker segments (residues 311-345 in CaMKII-α) and the regulatory segments (residues 281-310). The regulatory segments and linkers were modeled in arbitrary conformations, and located at variable distances from the surface of the dodecameric hub assembly. A 13 µs molecular dynamics trajectory was generated for this system. We also generated a 6 µs trajectory of the hub assembly without the linkers and regulatory segments. The lengths of the trajectories were limited by the availability of time on the Anton2 supercomputer.

In the simulation with the regulatory segments present, two of the twelve regulatory segments dock on the hub (Figure 4,5). Using the subunit notation shown in Figure 5A, these are the regulatory segments of subunits F and L, which dock at the interfaces between subunits B and L, and subunits F and H, respectively (Figure 4). The docking involves backbone hydrogen-bond formation between the calmodulin-binding element (residues 292-310) and the open edge of the β-sheet (residues 410-416) located at the interface (Figure 4S). The regulatory segment of subunit L docks within 0.5 µs (hereafter referred to as Docking 1), and that of subunit F docks at ∼3 µs (referred to as Docking 2) (Figure 5B).

**Figure 4.**
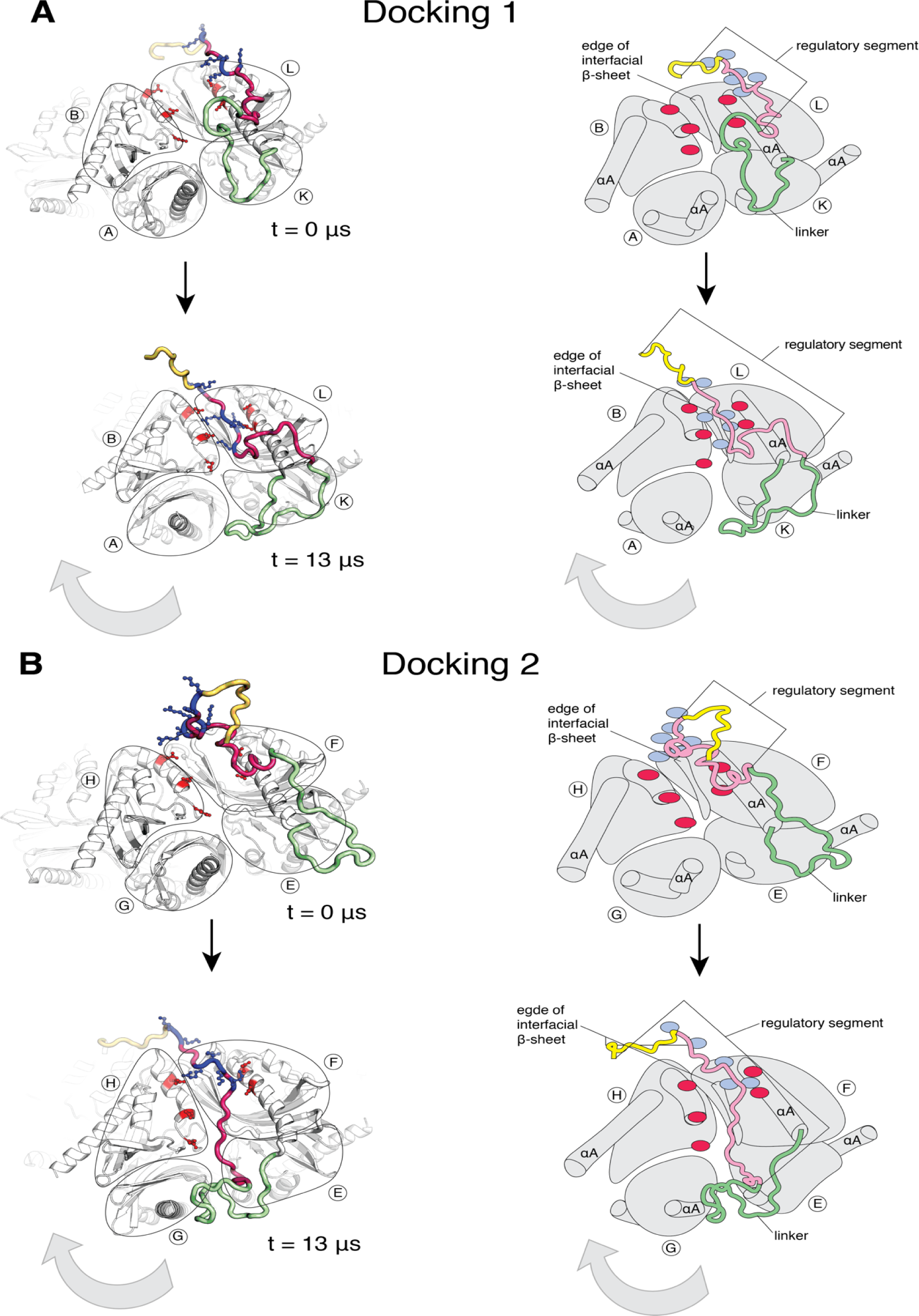
Calmodulin-binding elements from two of the twelve regulatory segments dock on the hub in the simulation of the dodecameric CaMKII-α hub, with considerable distortions at the interfaces where they bind. The color scheme for the regions of the subunits are the same as in Figure 1A. **(A)** Instantaneous snapshot from the start (top) and end (bottom) of the simulation showing interface where Docking 1 occurs. The calmodulin-binding element of subunit L docks at the interface between the hub domains of subunits B and L. The positively-charged residues of the regulatory segment (shown in blue) form interactions with the negatively-charged residues (shown in red) that line the interface between subunits B and L. The hub domain of subunit B rotates away from the hub domain of subunit L. **(B)** Instantaneous snapshot from the start (top) and end (bottom) of the simulation showing interface where Docking 2 occurs. The calmodulin-binding element of subunit F docks at the interface between the hub domains of subunits F and H. The positively-charged residues of the regulatory segment (shown in blue) form interactions with the negatively-charged residues (shown in red) that line the interface between subunits F and H. The hub domain of subunit H rotates away from the hub domain of subunit F.

**Figure 5.**
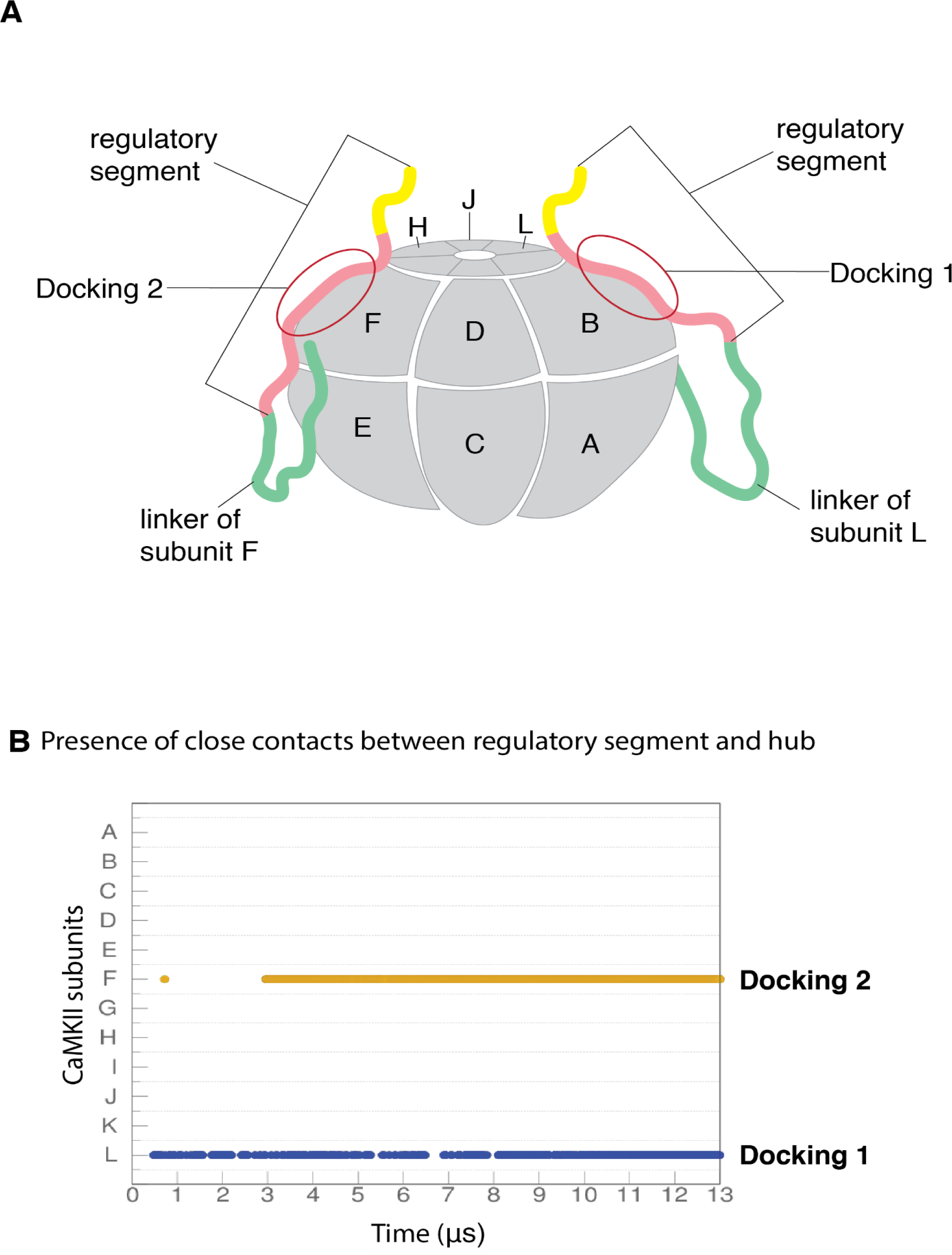
Docking of regulatory segments onto hub assembly. **(A)** A schematic representation of the hub assembly with the labeling convention used in this study. The two regulatory segments that dock on the hub assembly are shown, and colored using the same scheme as in Figure 1A. **(B)** Presence of interactions between residues 298-306 of the calmodulin-binding element of each subunits and the corresponding interfacial β-sheet (residues 410-416) over the course of the simulation. An interaction is considered to be present if at least two backbone hydrogen bonds (distance between donor and acceptor of less than or equal to 3Å) are formed between the calmodulin-binding element and the β-sheet.

Both docking events are initiated by interactions between positively-charged residues on the regulatory segments and negatively-charged residues that line the interfacial groove (Figure 5SB), although there are differences in the specific interactions that are made and in the register of the regulatory segments within the interfacial grooves. Backbone hydrogen bonds are then formed between the regulatory segment and the β-sheet at the interface, leading to the incorporation of this portion of the regulatory segment into the β-sheet, similar to the β-clip (Figure 4S) (Chao et al., 2011). The hydrophobic sidechains of the β-clip segment pack into the hydrophobic core of the subunits, further stabilizing docking by the regulatory segment. Unlike the electrostatic interactions, which are transient, the backbone hydrogen bonds are stable and persist for the duration of the simulation (Figure 4S).

The interfaces at which two dockings occur are located diametrically across from each other on the hub (Figure 5A) and docking is accompanied by a large conformational change at both interfaces (Figure 4A, B). We quantified this distortion by measuring the angle between the axes of the N-terminal α-helices (the αA helices) of two adjacent subunits (Figure 6A). The core structures of the individual subunits remain relatively unchanged over the course of the simulation (Figure 6SA), allowing us to use the angle between the axes of the αA helices as a measure of the overall rotation of one subunit at an interface with respect to the other. The initial value of the inter-helix angle between the αA helices of subunits B and L, which bracket the interface at which Docking 1 occurs, is ∼50°. These subunits begin to rotate away from each other when the trajectory is initiated, and the angle between helices αA has increased to ∼60° when Docking 1 is initiated (Figure 6C). At this point, there is a marked increase in the angle between the αA helices (to ∼80°), where it remains for the rest of the trajectory (Figure 6C).

**Figure 6.**
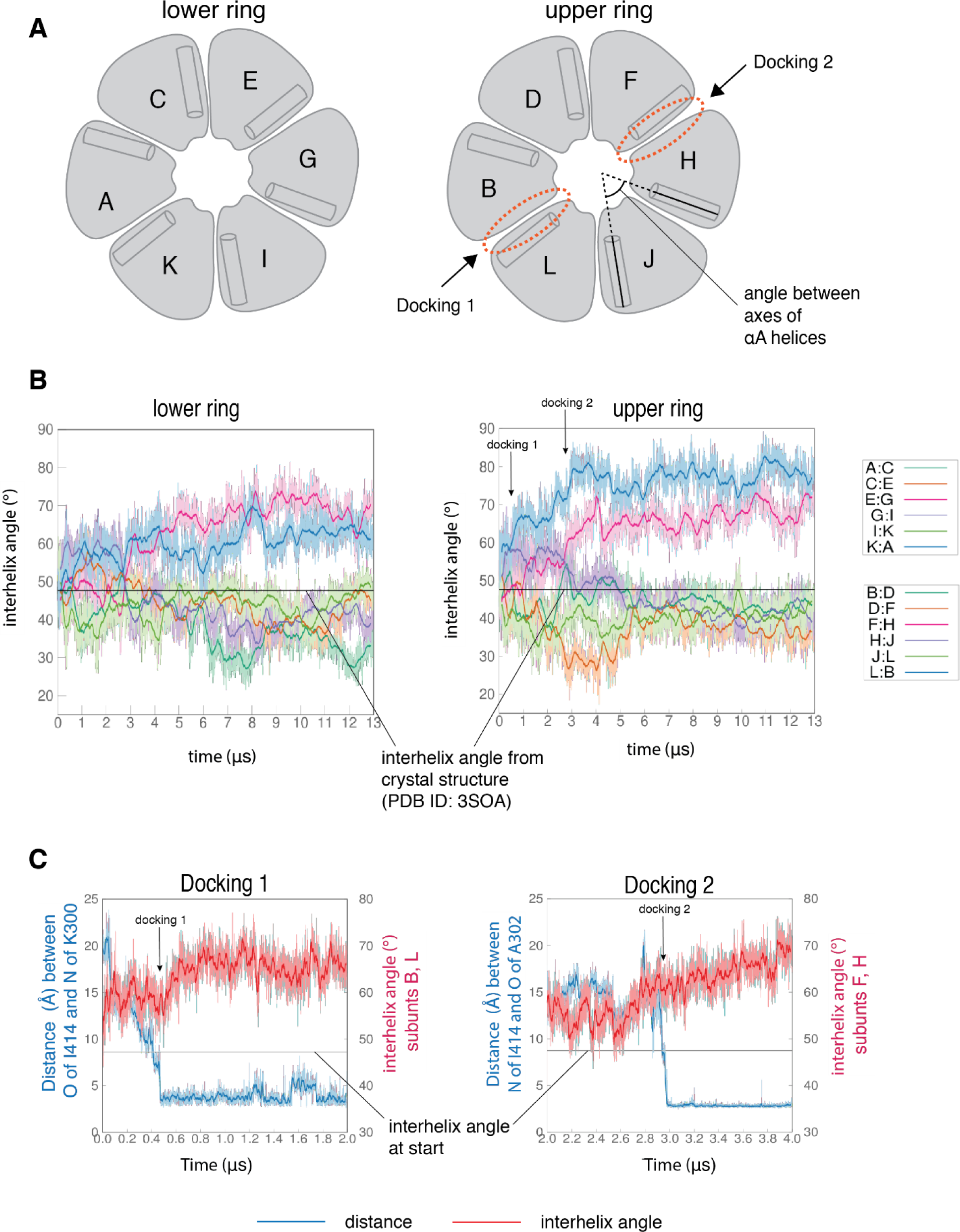
Interfaces at which docking occur are stabilized in an open form, while remaining interfaces close. **(A)** Schematic showing the naming scheme used to identify subunits. The interhelix angle between axes of αA helices are used to measure the rotation of each hub domain with respect to the adjacent hub domain. **(B)** Variation of the angle between the axes of the αA helices of adjacent subunits in the simulation with the regulatory segment. The darker traces are the time-averaged values of the interhelix angles calculated using a moving window of 240 ns, while the lighter shades are the actual distances. The two interfaces where regulatory segments dock between subunits L and B (colored blue), and between subunits F and H (colored pink) open and the remaining interfaces close. In the lower ring the interfaces mirror the behavior of the interfaces in the upper ring. The interhelix angle between the αA helices of adjacent subunits from the crystal structure of an autoinhibited, dodecameric holoenzyme (PDB ID - 3SOA) (Chao et al, 2011), which is perfectly symmetrical, is indicated by the horizontal black line. **(C)** Variation in interhelix angle between the axes of the αA helices of hub domains that flank the interfaces where the calmodulin-binding elements dock (shown in red), and the distance between a residue from the calmodulin-binding element and the interfacial β-sheet (shown in blue), over the course of the simulation. The darker traces are the time-averaged values, calculated using a moving window of 12 ns, while the lighter shades are the actual distances. The interhelix angle at the start of the simulation is indicated by a horizontal grey line.

Subunits F and H, which bracket the interface where Docking 2 occurs, also rotate away from each other from the start of the simulation (Figure 6C). The angle between the αA helices at this interface increases from an initial value of ∼50° to ∼60° by 3 µs, when Docking 2 is initiated, and then continues to increase over the rest of the simulation (Figure 6C). During the course of the simulation, the hub distorts from its normal circular shape to an oval shape, but it does not actually separate at any of the interfaces. This is because the large distortions at the two docking interfaces are compensated for by closure of the remaining interfaces (Figure 6B). This feature of the simulations suggests that the docking of additional segments to the hub is likely to result in a loss of the integrity of the hub assembly.

The rotation between the subunits that bracket the docking interfaces is much larger than observed in any crystal structures of the hub, including those with peptides bound at interfaces (Bhattacharyya et al., 2016; Chao et al., 2011). In a structure of the hub assembly of CaMKII from the sea anemone *Nematostella vectensis*, the N-terminal portion of the linker preceding the hub folds back to interact with the hub (Bhattacharyya et al., 2016). In this structure, as in the simulation, the linkers are incorporated as an additional strand of the interfacial β-sheet. The *N. vectensis* hub assembly is slightly cracked, and it assumes a spiral, lock-washer conformation (Bhattacharyya et al., 2016). The interaction between the linker and the *N. vectensis* hub involves acidic residues in the linker interacting with acidic residues in the hub. This interaction is likely to be an artefact of the low pH of crystallization and the consequent neutralization of acidic residues (Bhattacharyya et al., 2016). In contrast, the interaction between the regulatory segment and the hub seen in the simulation involves favorable interactions between oppositely charged sidechains.

### Regulatory segment docking appears to trap an intrinsic mode of distortion of the hub

It is striking that in the simulation a very similar distortion is seen at both docking sites, even though the register of the regulatory segment at one inter-subunit groove is offset by four residues with respect to the docking at the other inter-subunit groove (Figure 4S). This suggests that the distortion at the two docking sites reflects an intrinsic feature of the hub dynamics, one that does not depend on the details of the interaction with the peptide. In a 6 µs simulation of the hub without the regulatory segments, the hub assembly showed large structural excursions at interfaces between hub domains, including ones that are similar to those observed in the simulation of the hub with regulatory segments (Figure 6SB). Large displacements of subunits were also seen in a short 100 ns simulation of the hub reported previously (Bhattacharyya et al., 2016).

To better understand the intrinsic dynamics of the hub assembly we used normal mode analysis. We determined the normal modes of a dodecameric hub assembly (Chao et al., 2011), without the linkers, regulatory segments and the kinase domains, using the *ElNémo* server (Suhre and Sanejouand, 2004a, 2004b). This server uses the Elastic Network Model (Tirion, 1996) and the rotation–translation block method (Durand et al., 1994; Tama et al., 2000) to compute the normal modes.

To analyze the normal modes, we generated pairs of structures representing the sweep of each normal mode. Such pairs of structures were compared to the initial and final structures from the molecular dynamics trajectory. For each comparison, one of the two subunits at an interface was aligned between the pairs of structures, and the deviation in Cα positions of the adjacent subunit was considered. This gives us a measure of the differences in orientations of adjacent subunits between the normal mode displacement and the molecular dynamics trajectory.

We observed a striking similarity between the displacement seen in the molecular dynamics simulation and that seen in the lowest-frequency internal modes (Figure 7A,B). In these normal modes, just as in the molecular dynamics simulation, two of the inter-subunit interfaces open, and the other interfaces close. Thus, it appears that the coupled opening of two interfaces and the correlated closing of the others is an intrinsic property of the hub assembly. In the molecular dynamics simulation, it appears that the regulatory segments take advantage of the opening of the interfaces to form stable interactions with the interfaces and trap this distorted structure.

**Figure 7.**
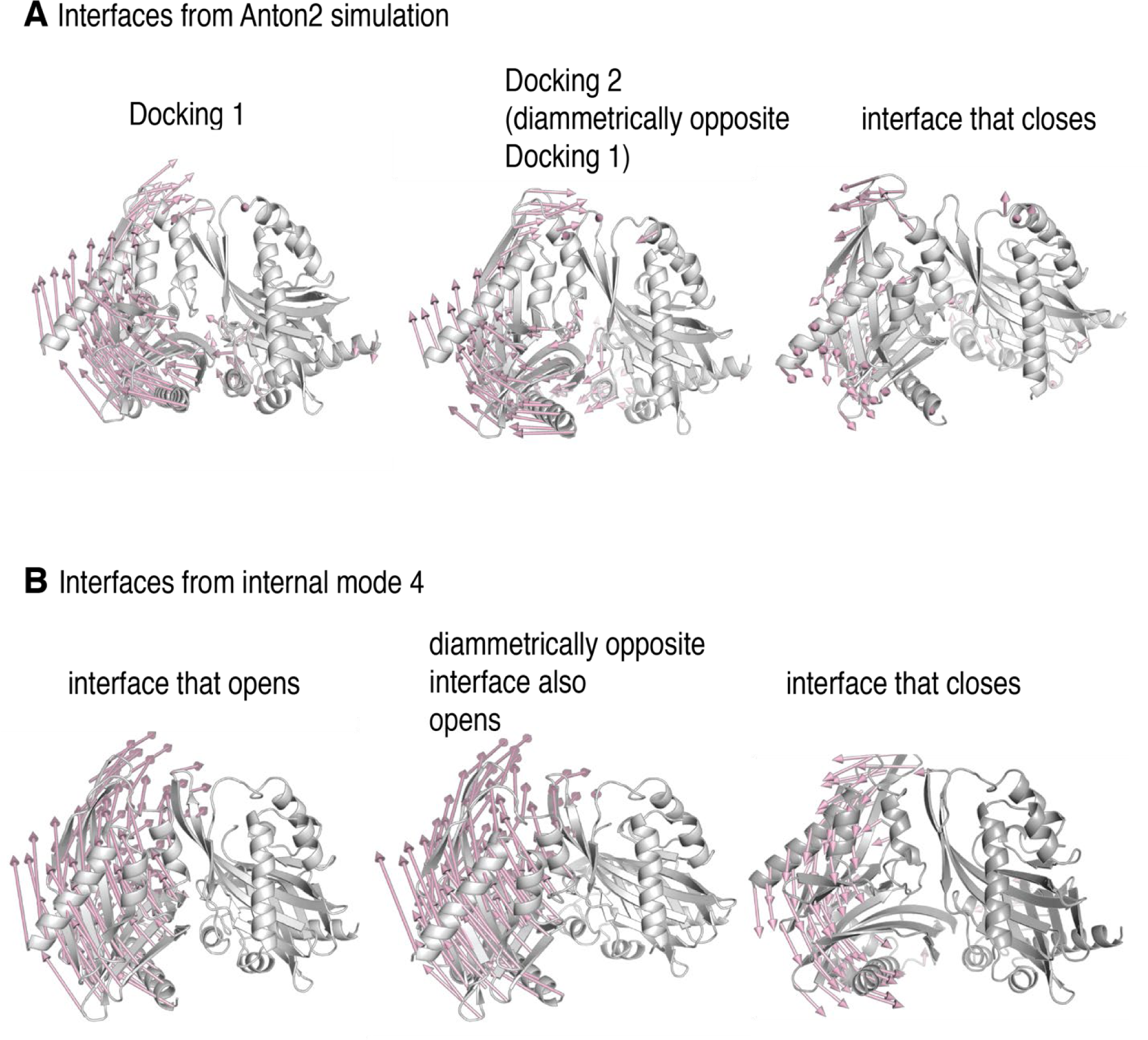
Docking traps the hub assembly in a conformation similar to one of its low frequency normal modes. **(A)** Conformational distortion in interfaces from the molecular dynamics simulation. The cartoon representations show the interfaces at the start of the simulation. Vectors (in pale pink) indicate the movement of the Cα-atoms of the hub domains to the left of the interfaces, with respect to the hub domains to the right of the interfaces, over the course the simulation. The interface at which Docking 1 (left) and Docking 2 (center) occur, open. One of the interfaces that closes is shown (right), and in this the hub domains to the left of the interface moves inwards, towards the interface. For clarity, only vectors from every other Cα-atom are shown. **(B)** Conformational change of interfaces in internal mode 4 from the normal mode analysis. The vectors (in pale pink) indicate the movement of the Cα-atoms of the hub domains to the left of the interfaces, with respect to the hub domains to the right of the interfaces, along the vector. Two of the interfaces (left and center) in internal mode 4 open, just as in the simulation, while the remainder (representative interface shown on right) close. For clarity, only vectors from every other Cα-atom are shown.

### Activation-induced destabilization of CaMKII-α holoenzymes expressed in mammalian cells

We studied the effect of activation on the stability of the intact CaMKII-α holoenzyme expressed in mammalian cells, using TIRF microscopy and single-molecule analysis. We developed a cell-based assay that allowed us to rapidly isolate CaMKII-α after cell lysis, while reducing the heterogeneity that can result from proteolysis or the aggregation that occurs during purification (Bhattacharyya et al., 2020). Briefly, CaMKII-α was tagged at the N-terminus with a biotinylation tag and mEGFP (Cormack et al., 1996; Zacharias et al., 2002) and overexpressed in HEK293T cells. The cells were lysed, and CaMKII-α was pulled down from the cell-lysate onto a streptavidin-coated glass coverslip (Figure 8A). CaMKII-α is visualized on the glass slide as spots of fluorescence intensity.

**Figure 8.**
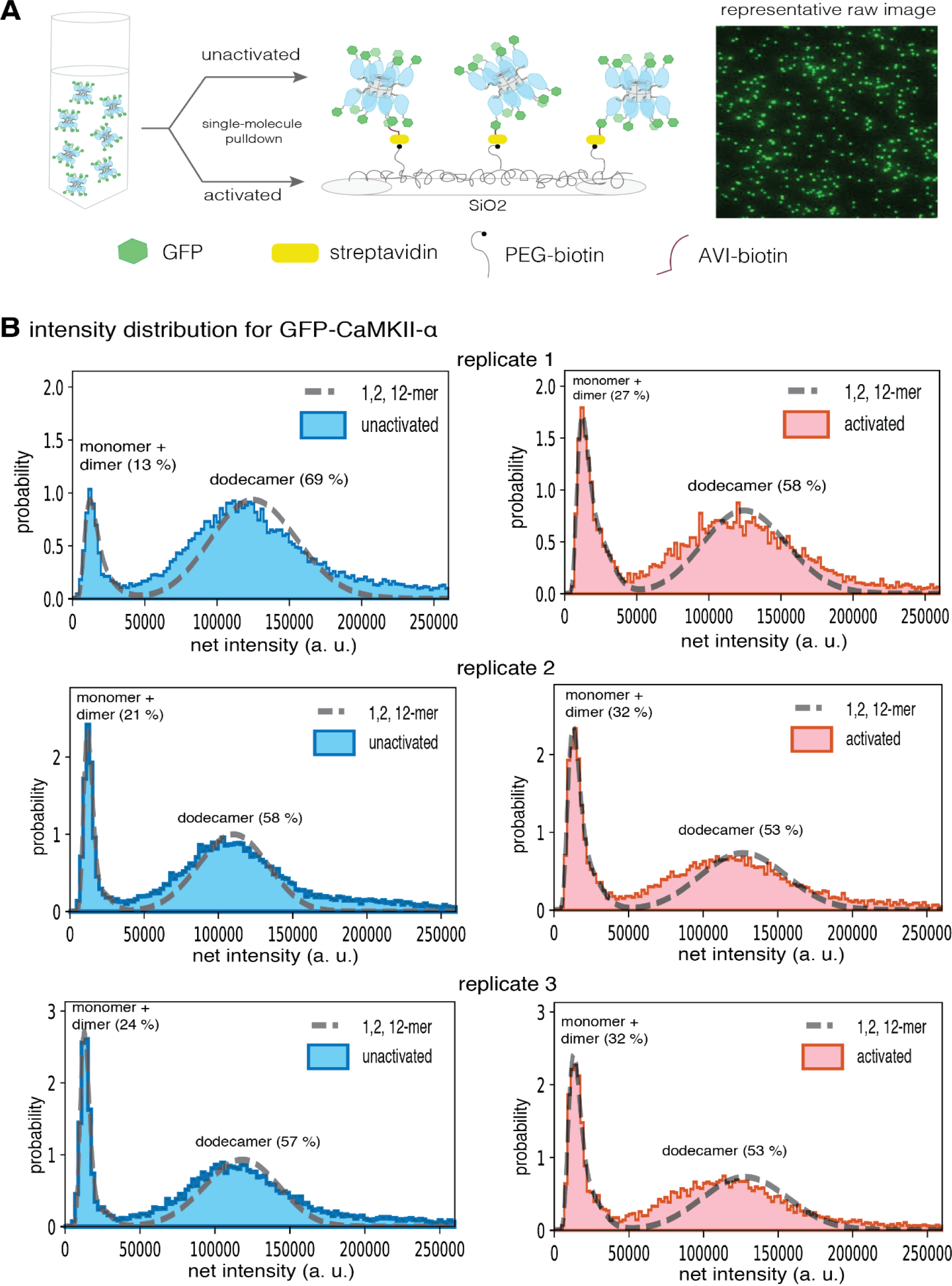
Activation destabilizes the CaMKII-α holoenzyme. **(A)** Schematic showing the experimental setup. Biotinylated mEGFP-CaMKII-α was overexpressed in HEK 293T cells, the cells were lysed and mEGFP-CaMKII-α from the diluted cell lysate was either first activated, or directly pulled down onto a glass coverslip, functionalized with streptavidin, for visualization at single-molecule resolution. The fluorescence intensity of each individual mEGFP-CaMKII-α is correlated with its oligomeric state. **(B)** Distribution of the intensities for three replicates of unactivated (left) and activated (right) mEGFP-CaMKII-α. Two major peaks are observed, with a peak at lower intensity, and a peak at higher intensity. There is a shoulder in the peak at lower intensity, and together these correspond to a mixture of monomers and dimers, with the dimer intensities occurring in the shoulder. The peak at higher intensities corresponds to intact dodecameric holoenzymes. The intensity data were fit to a mixture comprising of monomers, dimers and dodecamers (indicated by a dashed line) and the percentage of the smaller species and dodecameric species in the samples, as estimated from the fit are shown. Upon activation, the area of the peak at lower intensity rises, with a ∼1.3-2-fold increase across three replicates, indicating that upon activation the holoenzyme undergoes some disassembly to form dimers.

The distribution of fluorescence intensities for all the samples has two major peaks, one at lower intensities and one at higher intensities (Figures 8,9). Additionally, there is a shoulder in the lower intensity peaks in wild-type mEGFP-CaMKII-α (Figures 8). Step photobleaching analyses and a comparison with the intensity distribution of mEGFP-Hck, a monomeric protein kinase, indicate that the peaks with lower fluorescence intensity correspond to CaMKII-α monomers, while the shoulder corresponds to CaMKII-α dimers (Figure 8SB). Some percentage of GFP is dark and so a fraction of the spots with intensities corresponding to monomeric species actually correspond to dimeric species. For spots with higher fluorescence intensity in the CaMKII-α samples, photobleaching does not result in clearly resolvable steps (Figure 8SA).

**Figure 9.**
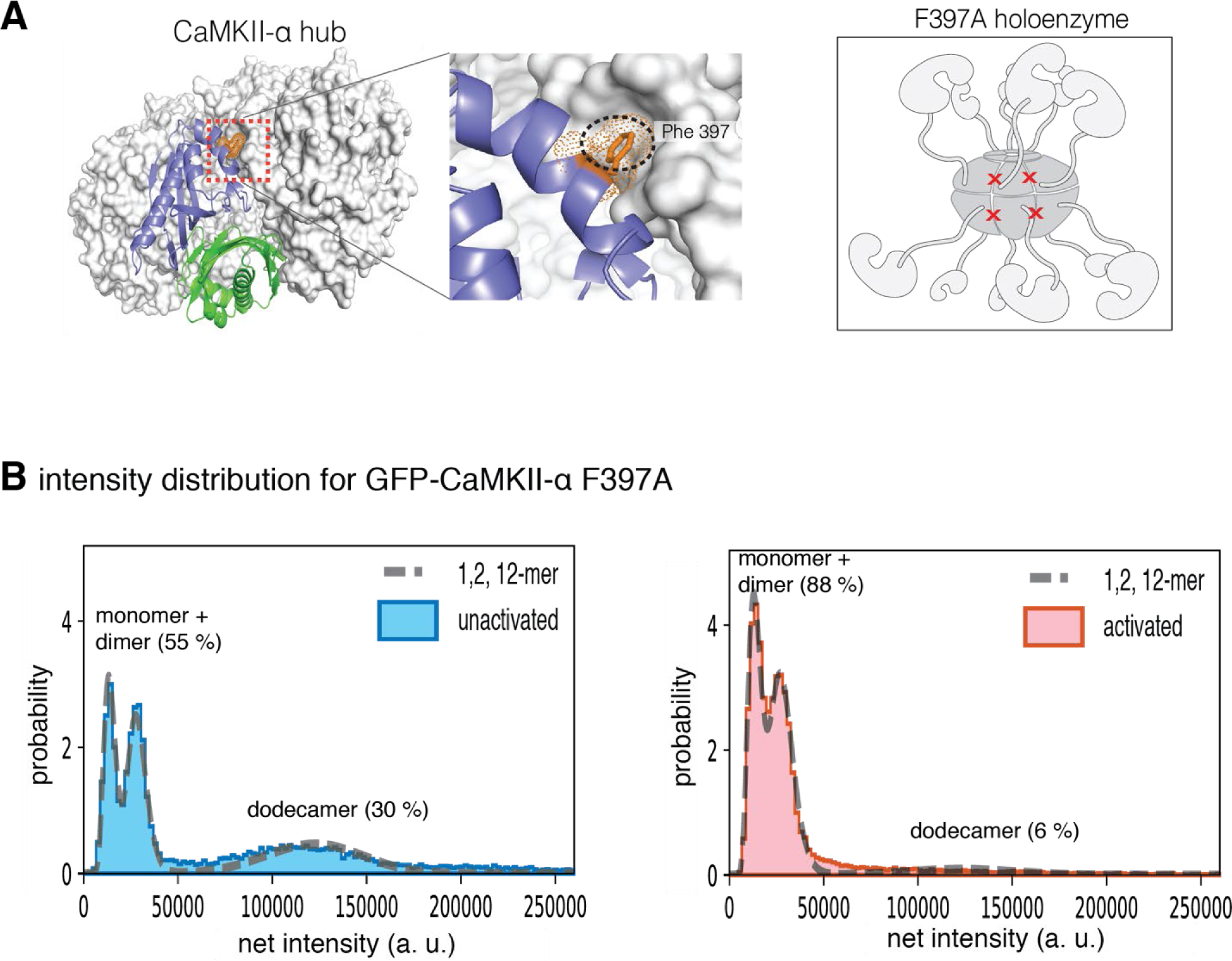
Intensity distribution of F397A mEGFP-CaMKII-α. **(A)** Crystal structure of human dodecameric CaMKII-α hub assembly (PDB ID – 5IG3) (Bhattacharyya et al, 2016) with the interface high-lighted. Replacement of Phe 397 at the hub interface with alanine weakens this interface. **(B)** Distribution of the intensities of unactivated (left) and activated (right) F397A mEGFP-CaMKII-α. At lower intensity, peaks corresponding to monomeric and dimeric species are observed. In the unactivated sample the peak at higher intensity, that corresponds to intact dodecameric holoenzyme is much smaller than in the wild-type sample. Upon activation this peak is no longer observed. The intensity data were fit to a mixture comprising of monomers, dimers and dodecamers (indicated by a dashed line) and the percentage of the smaller species and dodecameric species in the samples, as estimated from the fit are shown.

We estimated the number of subunits that comprise the peak at higher intensity by adapting the method described by Mutch et al (Mutch et al., 2007), where the position of the intensity peak for an oligomer with N subunits can be determined by convolving the monomeric (log-normal) distribution peak with itself N times. We also estimated the contribution of dark GFP and accounted for this in our calculation of the position of the N-oligomer peak (see Methods for more details). Approximately 30% of the GFP across all samples was found to be dark (Table 1B), consistent with previous findings (Ulbrich and Isacoff, 2007). Here, we fit the peak at lower intensity, extracted the parameters for the underlying monomeric log-normal distribution, and convolved it with itself twelve times, while accounting for dark EGFP, to predict the location and width of a peak comprising of dodecameric species. The predicted dodecamer intensity distribution corresponds well with the experimentally observed fluorescence intensity distribution of mEGFP-CaMKII-α (Figure 8B), indicating that this peak corresponds to predominantly dodecameric holoenzymes. From the fit, we were able to determine the percentage of smaller (monomeric and dimeric) species and larger, dodecameric species in the sample (Figure 8B, Table 1A). We have not accounted for the expected presence of a fraction of tetradecameric species. The ratio of dodecameric and tetradecameric species in assembled holoenzymes has been found to be variable, and in our analysis of CaMKII isolated from mammalian cells it appears to be small, consistent with recent analysis by electron microscopy (Myers et al., 2017).

**Table 1.** A distribution of oligomeric species

|  | **1** | **2** | **12** |
| --- | --- | --- | --- |
| Unactivated WT 1 | 0.103 | 0.028 | 0.693 |
| Activated WT 1 | 0.151 | 0.118 | 0.582 |
| Unactivated WT 2 | 0.169 | 0.037 | 0.583 |
| Activated WT 2 | 0.228 | 0.087 | 0.528 |
| Unactivated WT 3 | 0.196 | 0.042 | 0.574 |
| Activated WT 3 | 0.233 | 0.087 | 0.527 |
| Unactivated FA | 0 | 0.555 | 0.299 |
| Activated FA | 0.056 | 0.819 | 0.064 |

**B** estimated fraction of dark GFP
|  | **NUs** |
| --- | --- |
| Unactivated WT 1 | 0.282 |
| Activated WT 1 | 0.279 |
| Unactivated WT 2 | 0.328 |
| Activated WT 2 | 0.287 |
| Unactivated WT 3 | 0.296 |
| Activated WT 3 | 0.299 |
| Unactivated FA | 0.274 |
| Activated FA | 0.332 |

For unactivated mEGFP-CaMKII-α, the intensity distributions indicate the presence of a small population of monomers and dimers and a larger fraction of intact dodecameric holoenzymes (Figure 8B). We activated CaMKII-α in the diluted cell lysate with saturating amounts of Ca^2+^/CaM (5 µM) and ATP (10 mM), captured the activated CaMKII-α on a coverslip functionalized with streptavidin, and measured the distribution of fluorescence intensity of the imaged spots. These conditions have been shown previously to result in robust phosphorylation of CaMKII-α on Thr 286, with a lower amount of phosphorylation on Thr 305/Thr 306 (Bhattacharyya et al., 2020).

The fluorescence intensity distribution for CaMKII-α, along with the modeled distribution and the estimated percentages of the respective oligomeric species are shown in Figure 8B. There is a shift in the intensity distribution curve when compared to unactivated CaMKII-α, with the peak that corresponds to smaller (monomeric and dimeric) CaMKII-α showing a ∼1.3-2-fold increase in peak area, indicating that upon activation some of the intact holoenzyme undergoes disassembly to form smaller oligomeric species. The largest increase appears to be in the population of dimeric species (Table 1A), which is consistent with previous reports that the CaMKII holoenzyme is formed by the oligomerization of dimers (Bhattacharyya et al., 2016). The variation between the different replicates observed here is expected, as the release of dimers and monomers is a non-equilibrium process, with different amounts of smaller species present in the solution at the point of their capture on the glass.

The effect of activation upon CaMKII-α stability is amplified by the introduction of a destabilizing mutation at the hub interface (F397A) (Bhattacharyya et al., 2016) (Figure 9A). The distribution at lower intensities for this sample has two clearly resolvable peaks that correspond to a dimer population with 30 % dark GFP (Figure 9, Table 1A). The unactivated sample of this mutant form has a much larger percentage of smaller species when compared to any of the wild-type samples (Figure 9, 9S). Upon activation, the peak corresponding to intact holoenzymes is no longer observed (Figure 9, 9S), suggesting that the activation leads to complete destabilization of the CaMKII F397A variant holoenzyme.

## Conclusions

The unusual oligomeric organization of CaMKII, in which twelve or more kinase domains are arranged around a central hub, is coupled to a mechanism involving phosphorylation-mediated acquisition of constitutive activity that is also unique. When unphosphorylated, CaMKII is dependent on Ca^2+^/CaM for kinase activity. Once phosphorylated at Thr 286, CaMKII subunits acquire calmodulin-independent activity (autonomy). Thr 286 phosphorylation must necessarily occur in trans, with one kinase subunit phosphorylating another. Efficient Thr 286 phosphorylation occurs only within a holoenzyme (De Koninck and Schulman, 1998), with the enzyme-kinase and substrate-kinase being tethered to the same hub at high local concentrations. This feature makes subunit exchange potentially important for spreading the activation of CaMKII, since it provides a means for activated subunits to associate with subunits that have not yet been activated and thereby produce more activated subunits by trans-phosphorylation (Stratton et al., 2014).

The intriguing aspect of subunit exchange in CaMKII is that it is activation dependent. Most oligomeric proteins are capable of subunit exchange, provided that the dissociation rate constants are not too slow, and the subunits do not unfold when they are dissociated. Indeed, a CaMKII enzyme from choanoflagellates does not form stable oligomeric species, and it undergoes spontaneous subunit exchange (Bhattacharyya et al., 2016). In contrast, mammalian CaMKII-α is very stable as a dodecameric or tetradecameric oligomer and shows an insignificant degree of subunit exchange when not activated (Bhattacharyya et al., 2016; Stratton et al., 2014).

In this paper we have shown, using electrospray ionization mass spectrometry, that the stable assembly formed by the CaMKII-α hub is destabilized by the addition of peptides derived from the regulatory segment of CaMKII-α. Addition of regulatory segment peptides at 10 µM or higher concentration to samples of the hub leads to rapid loss of integrity of the hub, and the release of monomers, dimers, tetramers, hexamers and octamers. Molecular dynamics simulations indicate that individual subunits undergo considerable displacements with respect to adjacent subunits. These transient fluctuations lead to the interfacial grooves between subunits opening up, allowing regulatory segments to be captured. The simulations indicate that although two regulatory segments can dock on a closed hub, the binding of additional segments could disrupt the integrity of the hub. We have also shown, using a single-molecule TIRF assay, that activation of CaMKII-α expressed in mammalian cells results in destabilization of the holoenzymes.

We had suggested previously that there is a three-way competition for binding to the calmodulin-binding element between the kinase domain, Ca^2+^/CaM and the hub interface (Bhattacharyya et al., 2016). The calmodulin-binding element binds with nanomolar affinity to the kinase domain (Colbran et al., 1988), with picomolar affinity to Ca^2+^/CaM (Tse et al., 2007), and with micromolar affinity to the hub (Bhattacharyya et al., 2016). Upon activation, phosphorylation on Thr 286 prevents the calmodulin-binding element from binding to the kinase domains and phosphorylation on Thr 305/Thr 306 prevents interaction with Ca^2+^/CaM. Thus, if a CaMKII subunit undergoes phosphorylation at both Thr 286 and at Thr 305/Thr 306, then the regulatory segment might be released, and free to interact with the hub interface. The local concentration of the regulatory segments with respect to the hub is estimated to be in the millimolar range (Bhattacharyya et al., 2016). Given the observation that the addition of the regulatory segment peptides at micromolar concentrations in trans leads to hub disassembly, we conclude that the presentation of these elements in cis to the hub could lead to rapid destabilization of hub upon activation.

Subunit exchange in CaMKII has been proposed originally as a mechanism whereby the activation states of CaMKII could be maintained indefinitely, contributing to the storage of memories (Lisman, 1994). It is difficult to reconcile such a mechanism with the action of phosphatases, which are expected to reset the activation states of CaMKII (Bhattacharyya et al., 2020; Lee et al., 2009). Instead, we speculate that the destabilization of CaMKII holoenzymes upon activation is a mechanism that potentiates the effects of Ca^2+^ pulses under conditions when calmodulin is limiting, as is the case in neurons (Pepke et al., 2010; Persechini and Stemmer, 2002). Under these conditions, the release of dimers and other smaller oligomers can result in the spread of activated CaMKII species that can interact with and activate as yet unactivated holoenzymes. The precise mechanism by which this spread of activation occurs, and the role of subunit exchange in this process, awaits further study.

## Methods

### Preparation of plasmids

For the mass spectrometry analyses, the human CaMKII-α hub domain (Uniprot_ID: Q9UQM7, residue 345 - 475) was cloned into a pET-28 vector (Novagen) that was modified to contain a PreScission Protease (Pharmacia) site between the N-terminal 6-histidine tag and the coding sequence, before being expressed in *E. coli*. For the single-molecule analyses, the pEGFP-C1 vector was modified to contain a biotinylation sequence (Avitag, GLNDIFEAQKIEWHE) followed by a linker (GASGASGASGAS) at the N-terminus of mEGFP. Full-length human CaMKII-α (Uniprot_ID: Q9UQM7) was cloned into this vector backbone (Clontech), at the C-terminus of mEGFP, with a linker sequence (PreScission protease site: LEVLFQGP) separating the mEGFP tag from the coding sequence of CaMKII-α, before being overexpressed in mammalian cells (Bhattacharyya et al., 2020). This construct was then used as a template to produce a CaMKII-α variant (CaMKII-α-F397A). pET21a-BirA, which carries the *E. coli* biotin protein ligase sequence, was a gift from Alice Ting (Addgene plasmid # 20857). BirA was cloned into the pSNAP_f_ vector (New England Biolabs) after modifying the vector to remove the SNAP-tag. All constructs with insertions and deletions were made using standard protocols for Gibson assembly (New England Biolabs). All point mutants used were generated using standard Quikchange protocols (Agilent Technologies).

### Expression and purification of CaMKII-α hub

Human CaMKII-α hub domain was expressed in *E. coli* and purified as previously described (Bhattacharyya et al., 2016). Briefly, CaMKII-α hub protein expression was carried out in *E. coli* BL21 cells. Cells were induced by the addition of 1 mM isopropyl β-D-1-thiogalactopyranoside at an optical density of 0.7-0.8 and grown overnight at 18°C. Cell pellets were resuspended in Buffer A (25 mM Tris, pH 8.5, 150 mM potassium chloride, 1 mM DTT, 50 mM imidazole, and 10% glycerol) and lysed using a cell homogenizer. The cell lysate was filtered through a 1.1 micron glass fiber prefilter (ThermoFisher). The filtered lysate was loaded onto a 5 mL Ni-NTA column and eluted with Buffer B (25 mM Tris, pH 8.5, 150 mM potassium chloride, 1 mM DTT, 0.5 M imidazole, and 10% glycerol). The eluate was desalted using a HiPrep 26/10 desalting column into Buffer A with 10 mM imidazole, and cleaved with PreScission protease (overnight at 4°C). The cleaved samples were loaded onto a Ni-NTA column, the flow through was collected, concentrated and purified further using a Superose 6 gel filtration column equilibrated in gel filtration buffer (25 mM Tris, pH 8.0, 150 mM KCl, 1.0 mM tris(2-carboxyethyl)phosphine [TCEP] and 10% glycerol). Fractions with pure protein were pooled, concentrated and stored at −80°C. All purification steps were carried out at 4°C and all columns were purchased from GE Healthcare (Piscataway, NJ).

### Electrospray ionization mass spectrometry

Stock solutions of CAMKII-α hub were buffer-exchanged into 1 M ammonium acetate using a Bio-Spin column (Bio-Spin 6, Bio-Rad Laboratories, Inc., Hercules, CA). 10 mM stock solutions of different regulatory segment-derived peptides were prepared by dissolving the lyophilized powder in ddH_2_O. All peptides used in this study were purchased from Elim Biopharm, Hayward, CA. The hub and peptide were mixed such that final concentrations in the reaction mix contained 120 µM CaMKII-α hub (subunit concentration) and 1 µM to 1 mM of the peptide, with a constant ionic strength of ∼400 mM of ammonium acetate. This reaction mix was incubated for 5 minutes at room temperature before analysis by electrospray ionization mass spectrometry. As a control experiment, 120 µM of CaMKII-α hub in 400 mM ammonium acetate was also incubated for five minutes at room temperature and analyzed using electrospray ionization mass spectrometry.

Mass spectra were acquired on a Waters SYNAPT G2-Si mass spectrometer (Waters, Milford, MA, USA). Ions were formed from borosilicate emitters with a tip diameter of ∼500 nm that were pulled from capillaries (1.0 mm o.d./0.78 mm i.d., Sutter Instruments, Novato, CA, USA) using a Flaming/Brown micropipette puller (Model P-87, Sutter Instruments, Novato, CA, USA). The submicron emitters were chosen to help reduce salt adducts on the protein ions and to reduce the chemical noise (Susa et al., 2017). Nanoelectrospray ionization was initiated by applying a voltage of ∼1.0 kV to 1.2 kV onto a platinum wire (0.127 mm in diameter, Sigma, St. Louis, MO, USA) that is inserted inside the emitter and in contact with sample solutions. The emitter tips were placed ∼6 mm to 8 mm away from the entrance of the mass spectrometer.

To minimize collisional activation of the ions, a relatively low sample cone voltage (50 V) and low source temperature (80 °C) were used. The voltages of collisional induced dissociation cells (both Trap and Transfer cells) were set at 2 V, a value at which no gas-phase dissociation of the protein complex was observed. Mass spectral data were smoothed using the Savitsky-Golay smoothing algorithm, available via the Waters MassLynx software, with a smoothing window of 100 *m*/*z* (mass-to-charge ratio). For all experiments, the mass spectral results were continuously recorded as long as protein signals were observed. Mass spectra of the hub incubated with peptides were generated by averaging the scans for specific time segments after the initiation of electrospray ionization. These times correspond to the start of the mass spectrometry data acquisition and do not include the initial incubation time.

### System Preparation for Molecular Dynamics Simulations

For the long-timescale simulations, we built a dodecameric hub assembly by first building a model of a vertical dimer of hub domains connected to the regulatory segments by unstructured linkers. We ran short simulations of this structure while holding the hub domains fixed and allowing the linkers and regulatory segments to sample various conformations. We used instantaneous structures from these short simulations to build the dodecameric hub assembly with linkers and regulatory segments, to ensure that the simulation started with the regulatory segments in a variety of conformations at varying distances from the hub assembly.

For the short simulations of a vertical dimer, one vertical dimer of hub domains (residues 345-472) was taken from the crystal structure of the human CaMKII-α dodecamer (PDB ID-5IG3) (Bhattacharyya et al., 2016). Residues 336-344 at the N-terminus of the hub domain are missing in this structure and were modeled based on the structure of a tetradecameric CaMKII-α hub (PDB ID - 1HKX (Hoelz et al., 2003). The C-terminus of the hub domain (resides 473-478) and the linker (residues 314-335) were built in irregular conformations using Coot (Emsley and Cowtan, 2004) and PyMOL (Schrödinger, LLC., 2015), via the SBGrid Consortium (Morin et al., 2013). The N-terminal (residues 281-300) and C-terminal regions (residues 308-313) of the regulatory segment were modeled based on the structure of a CaMKII-calmodulin complex (PDB ID - 2WEL) (Rellos et al., 2010). Residues 301-307, which comprise the β-clip segment of the calmodulin-binding element, form a helix when bound calmodulin (Rellos et al., 2010). In the absence of calmodulin, this region is expected to unwind from a helical architecture, and so the starting structure for this region was modeled based on the structure of an autoinhibited CaMKII, where this portion of the helix is unwound (PDB ID - 2VN9) (Rellos et al., 2010). The N-termini of the individual subunits were capped with acetyl groups, because they represent truncations of the actual structure, whereas the C-termini were left uncapped.

The protein was solvated, and ions were added such that the ionic strength was 0.15 M, using VMD (Humphrey et al., 1996). The energy of this system was minimized for 1000 steps, first while holding the protein atoms fixed, then for 1000 steps while allowing all the atoms to move. It was then equilibrated by molecular dynamics at constant pressure (1 atm) and constant temperature (300 K) for 2 ns, then equilibrated at constant volume and constant temperature (300 K), to ensure that the pressure is maintained stably, for 1 ns. We generated two short trajectories (each ∼60 ns) at constant pressure (1 atm) and constant temperature (300 K), while holding hub domains (residues 336-478) fixed, and allowing the linkers and regulatory segments to explore various conformations.

Instantaneous structures from these simulations, with the linker and regulatory segment in arbitrary conformations, were then used to build a dodecameric assembly comprising of the hub, linker and regulatory segments, by aligning a pair of vertical hub domain dimers onto the vertical hub domain dimers of the human dodecameric hub assembly (PDB ID – 5IG3) (Bhattacharyya et al., 2016). This system was then solvated and ions were added so that the final ionic strength was 0.15 M using VMD (Humphrey et al., 1996). The final system had a cubic cell with edge dimensions of 193 Å.

A second system comprising of the dodecamer hub assembly only, without the linker and regulatory segments was built from the system described above, by removing residues 281-335 from each subunit. This system was solvated with TIP3P water and ions were added so that the final ionic strength was 0.15 M using VMD (Humphrey et al., 1996). The final system had a cubic cell with edge dimensions of 193 Å.

The energies of both systems were minimized first while holding the protein fixed, then while allowing all the atoms to move. The two systems were then equilibrated at constant pressure (1 atm) and temperature (300 K) for 5 ns. The NAMD package was used to run the minimization and equilibration simulations (Phillips et al., 2005) with the CHARMM36 force field (Best et al., 2012). The velocity Verlet algorithm was used to calculate the trajectories of the atoms. A time step of 2 fs was used. Particle Mesh Ewald was used to calculate long-range electrostatic interactions, with a maximum space of 1 Å between grid points (Darden et al., 1993). Long-range electrostatics were updated at every time step. Van der Waal’s interactions were truncated at 12 Å. Hydrogen atoms bonded to the heavy atoms were constrained using the ShakeH algorithm (Ryckaert et al., 1977). Temperature was controlled with Langevin dynamics with a damping coefficient of 4/ps, applied only to non-hydrogen atoms. Pressure was controlled by the Nose-Hoover method with the Langevin piston, with an oscillation period of 200 fs and a damping time scale of 50 fs (Feller et al., 1995; Martyna et al., 1994).

### Anton Simulation Protocols

Production runs for both systems were carried out on Anton2 (Shaw et al., 2014). The restart files from the end of the equilibration stage were converted to the Desmond format (Bowers et al., 2006) using the VMD (Humphrey et al., 1996). The CHARMM param36 force field (Best et al., 2012) was used through the Viparr utility of Anton2 (Shaw et al., 2014). A time step of 2 fs was used. The multigrator integration method (Lippert et al., 2013) was used to generate the trajectories while using the Nose-Hoover thermostat to maintain temperature at 300 K and the MTK barostat to maintain pressure maintain pressure at 1 atm (Lippert et al., 2013). Coordinates were saved every 0.24 ns. The coordinate trajectories were converted to the NAMD DCD format using VMD (Humphrey et al., 1996).

The angle between the αA helices (residues 341-363) of adjacent subunits were calculated using the CHARMM package (Brooks et al., 2009). The r. m. s. deviation for each hub domain over the course of the simulation of the dodecamer with the regulatory segment and linkers present was calculated with the AmberTools18 package (Case et al., n.d.), using residues 340-470 of the hub domain while fitting to the structure of the hub at the start of the simulation.

### Normal Mode Analysis

The hub assembly (residues 340:470) from the structure of the dodecameric autoinhibited CaMKII-α (PDB ID – 3SOA) was used for normal mode analysis. The *elNém*o server was used for the analysis with a minimum and maximum perturbation of 400, a step-size of 16 and a residue-block size of 8 (Suhre and Sanejouand, 2004a, 2004b). The displacement vectors were displayed using the modevectors script in PyMOL (Schrödinger, LLC., 2015).

### Tissue culture and DNA transfection

HEK 293T cells (UC Berkeley cell culture facility) were grown in DMEM (Dulbecco’s Modified Eagle Medium + GlutaMaX, ThermoFisher) supplemented with 10% fetal bovine serum (FBS), 1X antibiotic–antimycotic reagent (ThermoFisher) and 20 mM HEPES buffer and maintained at 37°C under 5% CO_2_. CaMKII-α variants were transiently transfected using the standard calcium phosphate protocol. Briefly, calcium phosphate/DNA coprecipitate was prepared by combining CaMKII-α plasmids with 5 μg of empty pcDNA3.1 vector and 1 μg of BirA plasmid.

The mixture was then diluted with 10X ddH_2_O, and CaCl_2_ was added such that the final CaCl_2_ concentration is 250 mM. This mixture was incubated for 15 minutes. One volume of this 2X calcium phosphate/DNA solution was added to an equal volume of 2X HEPES-buffered saline (HBS) (50 mM HEPES, 280 mM NaCl, 1.5 mM Na_2_HPO_4_, pH 7.1) and the solution was mixed thoroughly by reverse pipetting. This mixture was then added to HEK 293T cells and the cells were allowed to express the protein for 18-20 hours, before the protein was harvested.

### Preparation of flow cells for single-molecule microscopy

All single-molecule experiments were performed in flow chambers (sticky-Slide VI 0.4, Ibidi) that were assembled with functionalized glass substrates. The glass substrates (coverslips, Ibidi) were first cleaned using 2% Hellmanex III solution (Hellma Analytics) for 30 min, followed by a 30 min sonication in 1:1 mixture (vol/vol) of isopropanol:water. The glass substrates were then dried with nitrogen and cleaned for another 5 min in a plasma cleaner (Harrick Plasma PDC-32 G). These cleaned glass substrates were used to assemble the flow chambers immediately after plasma cleaning. After assembly, the glass substrates were treated with a mixture of Poly-L-lysine PEG and PEG-Biotin (1000:1, both at 1 mg/mL) for 30 min (SuSoS). The glass substrates were then washed with 2 mL of phosphate-buffered saline (PBS). Streptavidin was added to these glass substrates at a final concentration of 0.1 mg/mL and incubated for 30 min. Following incubation, excess streptavidin was washed away using 2 mL of PBS and these assembled flow chambers were used for all our single-molecule experiments.

### Cell lysis and pulldown of biotinylated CaMKII-α in flow chambers

The co-expression of the *E. coli* biotin ligase, BirA, with the CaMKII-α variants bearing an N-terminal Avitag, results in the biotinylation of the Avitag in HEK 293T cells. After harvesting, the cells were lysed in a lysis buffer (25 mM Tris at pH 8, 150 mM KCl, 1.5 mM TCEP-HCl, 1% protease inhibitor cocktail (P8340, Sigma)), 0.5% phosphatase inhibitor cocktails 2 (P0044, Sigma) and 3 (P5726, Sigma), 50 mM NaF, 15 μg/ml benzamidine, 0.1 mM phenylmethanesulfonyl fluoride and 1% NP-40 (ThermoFisher).

The cell lysate was diluted in the gel filtration buffer (same recipe as above, but without glycerol). mEGFP-CaMKII-α in the diluted cell lysate was activated by incubating the lysate with an activation buffer containing 5 μM CaM, 100 μM CaCl_2_, 10 mM ATP and 20 mM MgCl_2_ (final concentrations in the reaction) for 60 minutes at room temperature. For the unactivated sample, the 60-minute incubation was carried out in gel filtration buffer without the addition of any components of the activation buffer. For both the activated and unactivated samples, 100 μL of the diluted cell lysate was added to a well in the flow chamber. After incubation for 1 minute, the diluted cell lysate was washed out with 1 mL of PBS. During this incubation, the biotinylated mEGFP-CaMKII-α variants were immobilized on the surface of the functionalized glass substrates, via the streptavidin-biotin interaction.

### Single-molecule Total Internal Reflection Fluorescence (TIRF) microscopy

Single-particle total internal reflection fluorescence (TIRF) images were acquired on a Nikon Eclipse Ti-inverted microscope equipped with a Nikon 100x 1.49 numerical aperture oil-immersion TIRF objective, a TIRF illuminator, a Perfect Focus system, and a motorized stage. Images were recorded using an Andor iXon electron-multiplying CCD camera. The sample was illuminated using the LU-N4 laser unit (Nikon) with solid-state lasers for the channels emitting at wavelengths of 488 nm, 561 nm and 640 nm. Lasers were controlled using a built-in acousto-optic tunable filter. A 405/488/561/638 nm Quad TIRF filter set (Chroma Technology Corp.) was used along with supplementary emission filters of 525/50m, 600/50m, 700/75m for the 488 nm, 561 nm and 640 nm channels, respectively. Image acquisition was performed with the automated change of illumination and filter sets, at 75 different positions from an initial reference frame, so as to capture multiple non-overlapping images, using the Nikon NIS-Elements software. mEGFP-CaMKII-α was imaged by illuminating the 488 nm laser set to 5.2 mW. Images for all the mEGFP-CaMKII-α variants were acquired using an exposure time of 80 ms, while keeping the laser power unchanged over all conditions.

To correct for the baseline clamp and dark current, a series of dark images were collected under the same exposure time of 80 ms as the experimental sample. Uneven illumination and fringe interference effects in the intensity distribution (Mattheyses et al., 2010) were corrected for by measuring the field illumination with a solubilized fluorescein sample. The dark image was subtracted from both the sample data and field illumination control, then the sample data were divided by a normalized mean field control to acquire appropriate intensities.

For photobleaching experiments, the GFP-fluorescence signal was recorded in a stream acquisition mode, with an exposure time of 80 ms, under the same conditions as described above. As a control, the intensities and photobleaching traces of mEGFP-Hck, a monomeric protein kinase, were also collected under the same conditions.

### Analyses of single-molecule TIRF data

Individual single particles of mEGFP-CaMKII-α were detected and localized using the single particle tracking plugin TrackMate in ImageJ (Jaqaman et al., 2008). The particles were localized with the Laplacian of Gaussian detector with an initial diameter set to 6 pixels. A threshold value of 300 was used to exclude noisy, low-intensity spots. To eliminate the effects of variation in resolution at the edges of the field due to heterogenous TIRF illumination, only particles within a central area of 400 x 400 pixel^2^ were included in the calculation. The intensity distributions for single particles of mEGFP-CaMKII-α (488 nm) were determined using custom in-house software written in Python. The total intensity values acquired from TrackMate for the 488-channel were adjusted by subtracting the local median background intensity for each spot (scaled by the spot area) to produce an intensity histogram for all the mEGFP-CaMKII-α spots.

For the analysis of the GFP step-photobleaching data, the photobleaching traces for each spot were built by plotting the maximum intensities of mEGFP-CaMKII-α as a function of time with MATLAB. The number of single photobleaching events were counted manually by inspecting the photobleaching trace for every spot. Single-step and two-step bleaching traces were clearly identified, but multistep photobleaching traces exhibited relatively unclear, distinct bleaching events. We therefore categorized the bleaching traces into single-step, two-step, and multi-step photobleaching. A similar analysis was used for mEGFP-Hck, which exhibited only single step traces.

### Determination the composition of oligomeric species in the intensity distributions

The intensity distribution of monomeric GFP species is lognormal when detected on an EMCCD camera, and is given by

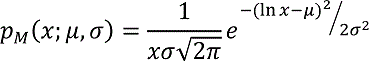

where *x* is the integrated intensity of a spot, µ is the mean of the underlying normal variable and σ is its standard deviation. Given a monomer distribution, the intensity distribution for a species with N subunits can be determined by convolving the intensity distribution of the monomer with itself N times (Mutch et al., 2007). An oligomer may have some dark GFP leading to a different intensity distribution than expected, and the dark fraction (ν) can be used to predict the extent to which this occurs. This was done by sampling *p_M_* finely, converting it to a probability mass function (pmf) and setting

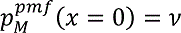

Subsequent convolutions of this modified probability mass function carry the effect of the dark GFP. The probability mass function was then reconverted back to a density function.

We estimated the dark fraction (ν) by auto-convolving the monomeric distribution 12 times and determining value of ν that yields the best alignment of the predicted dodecameric distribution with the high-intensity peak in the experimental data. These gave values of ν ∼ 0.3 (Table 1B). While this method of estimating ν will center the predicted peak with the data, it has no bearing on its width, which is governed by the underlying lognormal function. Based on these results, a dark fraction of 0.3 was used for all samples.

The following procedure was used to create the final mixture model in Figures 8 and 9. First the low-intensity peak was fit to a mixture of monomeric and dimeric species with a fixed dark fraction (ν) of 0.3, which yielded parameters σ and μ for the underlying monomeric lognormal distribution. This log-normal distribution was convolved with itself twelve times, also with a fixed dark rate of 0.3, to gives three final distributions: monomer, dimer, and dodecamer. The observed data were then fit to a mixture of these three populations.

## Acknowledgements

We thank Howard Schulman for helpful discussions. BX and ERW are supported by the National Science Foundation Division of Chemistry under grant number CHE-1609866. The authors also thank CALSOLV for funding. MB thanks NIGMS (K99 GM 126145) for funding. Preliminary simulations were run on Comet, at the San Diego Supercomputing Center, via Extreme Science and Engineering Discovery Environment (XSEDE), which is supported by National Science Foundation grant number ACI-1548562. Long-timescale simulations were carried out on Anton2. Anton 2 computer time was provided by the Pittsburgh Supercomputing Center (PSC) through Grant R01GM116961 from the National Institutes of Health. The Anton 2 machine at PSC was generously made available by D.E. Shaw Research.

---

**Figure 2 Supplement.**
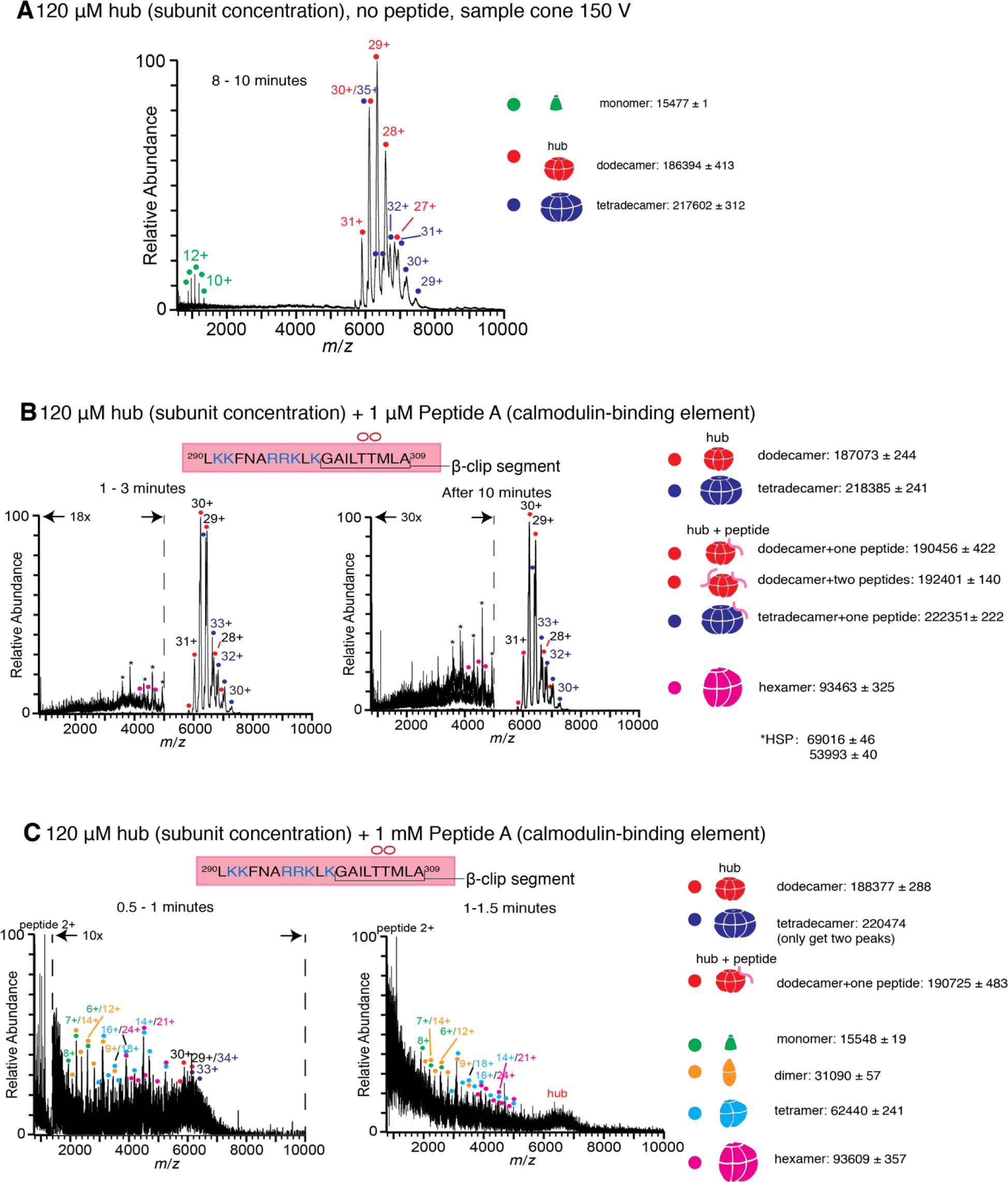
**(A)** Mass spectrum of the isolated hub with a sample cone voltage of 150 V. Monomeric species of the hub assembly are observed at this high sample cone voltage. **(B)** Incubation of the hub assembly with 1 µM of Peptide A shows no disassembly, even up to 10 minutes after ionization, and only dodecamers and tetradecamers are observed. **(C)** Incubation of the hub assembly with 1 mM of Peptide A shows significant protein/peptide aggregation. In all the spectra of the hub incubated with peptides, species corresponding to dodecameric or tetradecameric hub assemblies with one or two peptides bound are present, even when no hub disassembly is observed.

**Figure 3 Supplement.**
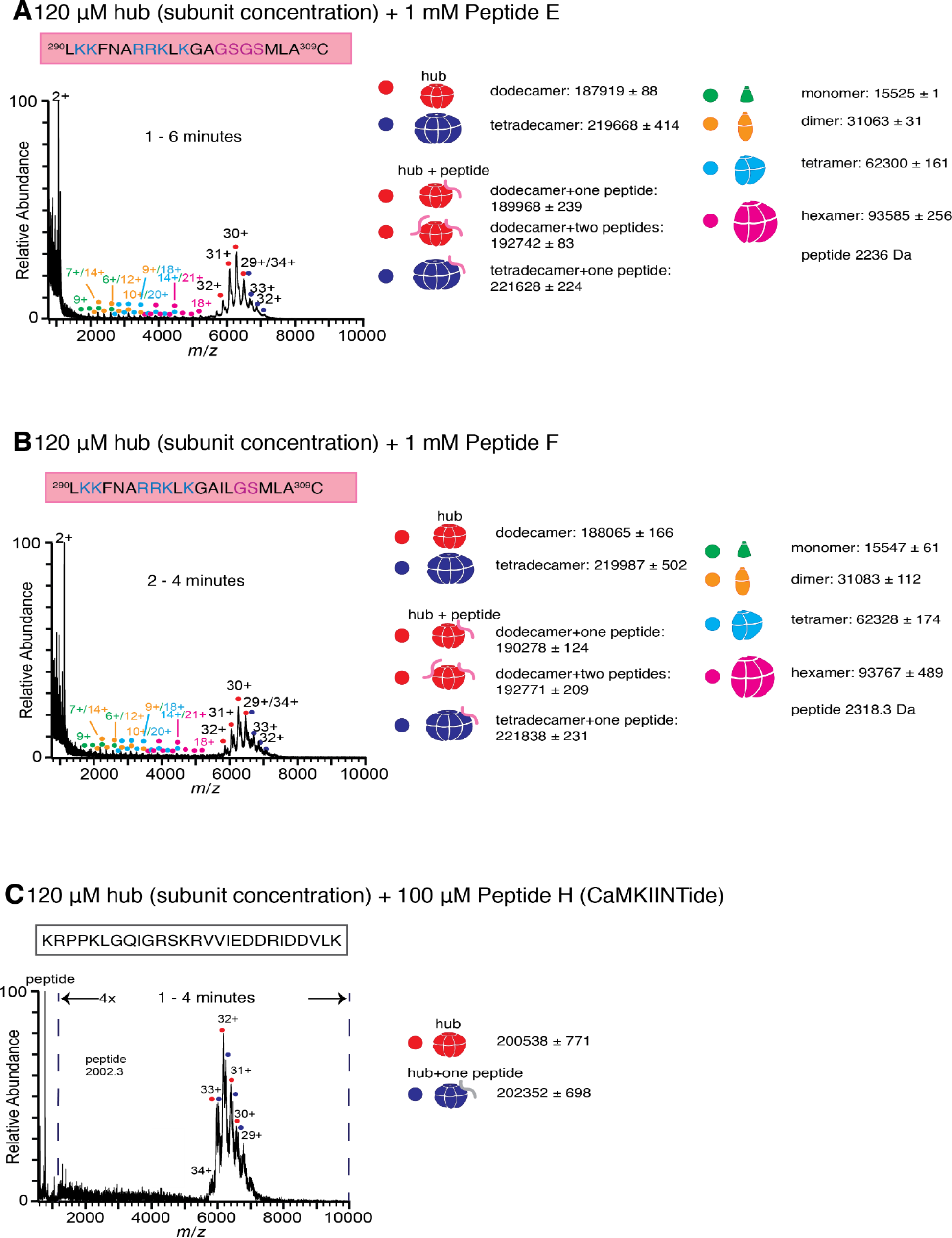
**(A)** Incubation of the hub assembly with 1 mM of Peptide E, in which a portion of the β-clip segment of Peptide A is replaced with glycine and serine residues (colored purple) results in the release of some smaller oligomeric species up to 6 minutes after ionization. **(B)** Incubation of the hub assembly with 1 mM of Peptide F, in which a portion of the β-clip segment of Peptide A is replaced with glycine and serine residues (colored purple) results in the release of some smaller oligomeric species up to 4 minutes after ionization. **(C)** Incubation of the hub assembly with 100 µM of Peptide H (CaMKIINTide), an inhibitor that binds to the substrate-recognition site of the kinase domain of CaMKII, does not result in the release of smaller oligomeric species up to 4 minutes after ionization. The CAMKII dodecamers and tetradecamers are not resolved in this mass spectra due to the soft instrument source conditions used, and peak broadening that resulted from nonvolatile salts present in the peptide solution.

**Figure 4 Supplement.**
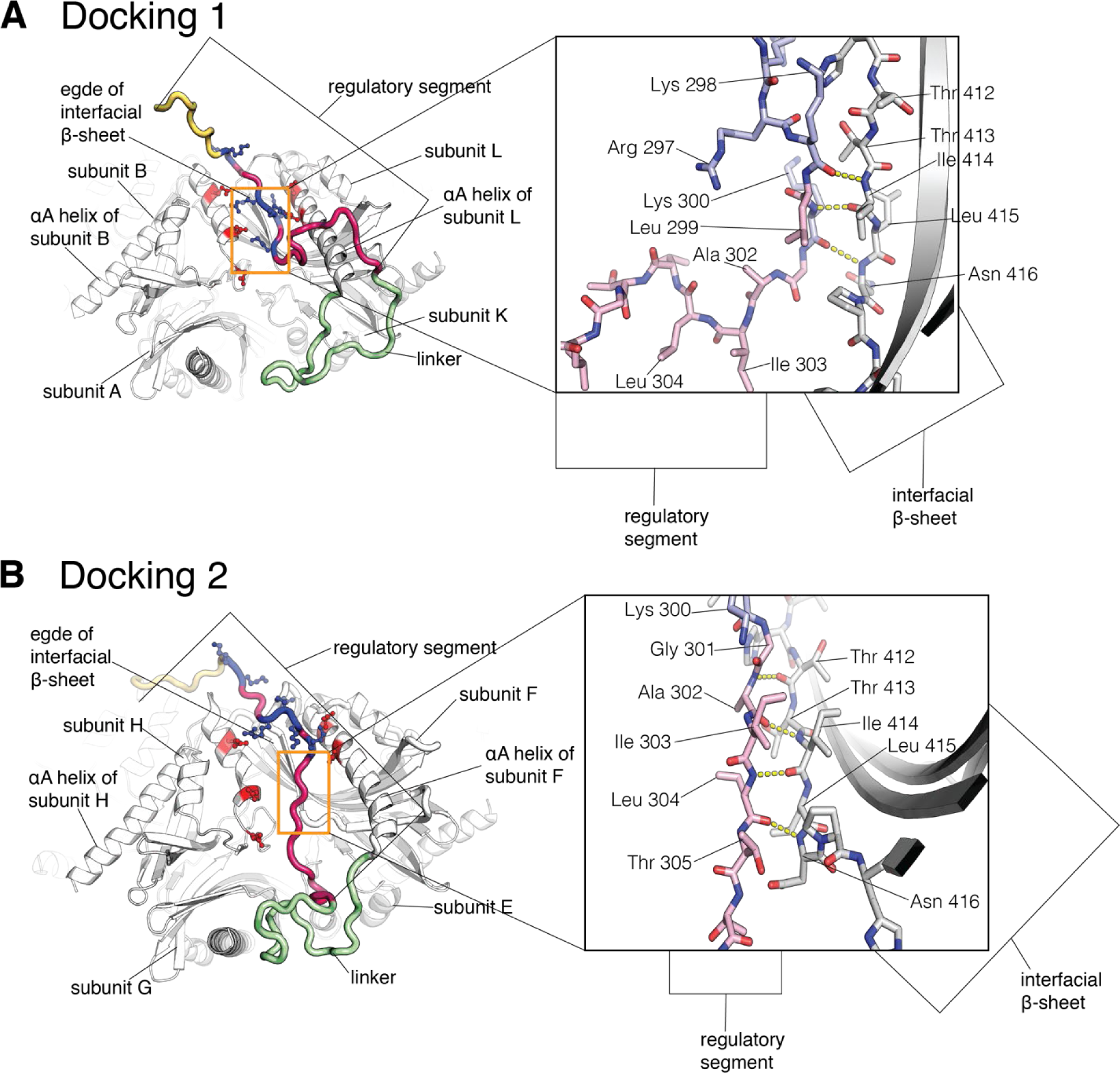
Close-up of two docking events in the simulation. The calmodulin-binding element is colored pale pink, with the positively charged residues colored blue. The interfacial β-sheet is shown in white. **(A)** Close-up of Docking 1 showing backbone interactions between residues 298-300 of the calmodulin-binding element of subunit L, and residues 412-416 of the edge of the β-sheet of subunit L. **(B)** Close-up of Docking 2 showing backbone interactions between residues 301-305 of the calmodulin-binding element of subunit F, and residues 412-416 of the edge of the β-sheet of subunit F.

**Figure 5 Supplement.**
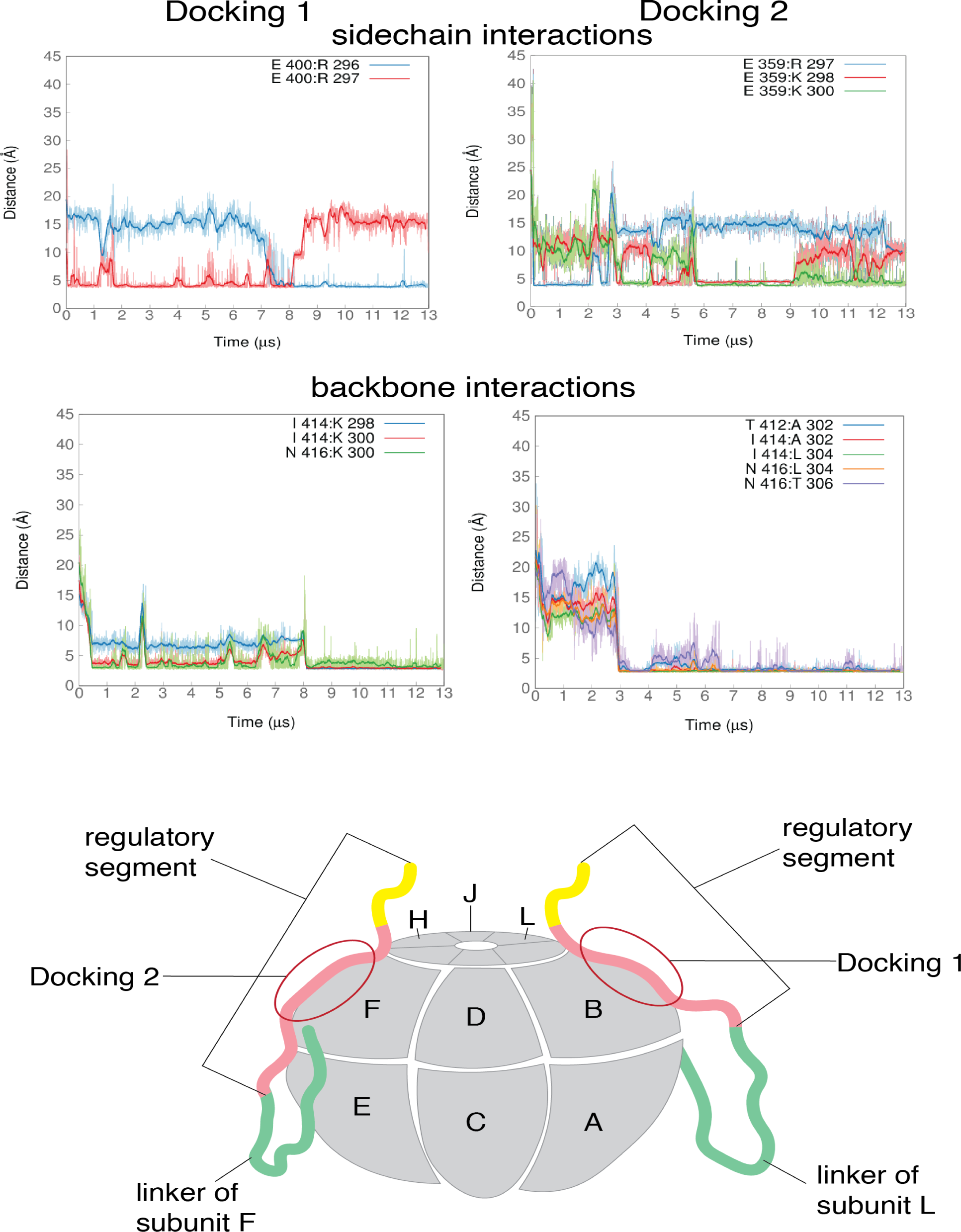
Sidechain interactions are formed soon after the start of the simulation and are not stable over the course of the simulations, while the backbone interactions are stable once they are formed. A representative sample of distances between residues of the calmodulin-binding element and residues at the interface of the hub assembly, from the two docking events are shown, with Docking 1 on the left and Docking 2 on the right. The darker traces are the time-averaged values of the distances calculated using a moving window of 470 ns while the lighter shades are the actual distances. The two figures on top show distances between positively-charged residues from the calmodulin-binding element and negatively-charged residues that line the interface between two subunits. The terminal carbon atoms of the sidechains were used to calculate distances. The two figures at the bottom show distances between the N and O atoms from residues of the backbone of the calmodulin-binding element and the backbone of the interfacial β-sheet.

**Figure 6 Supplement.**
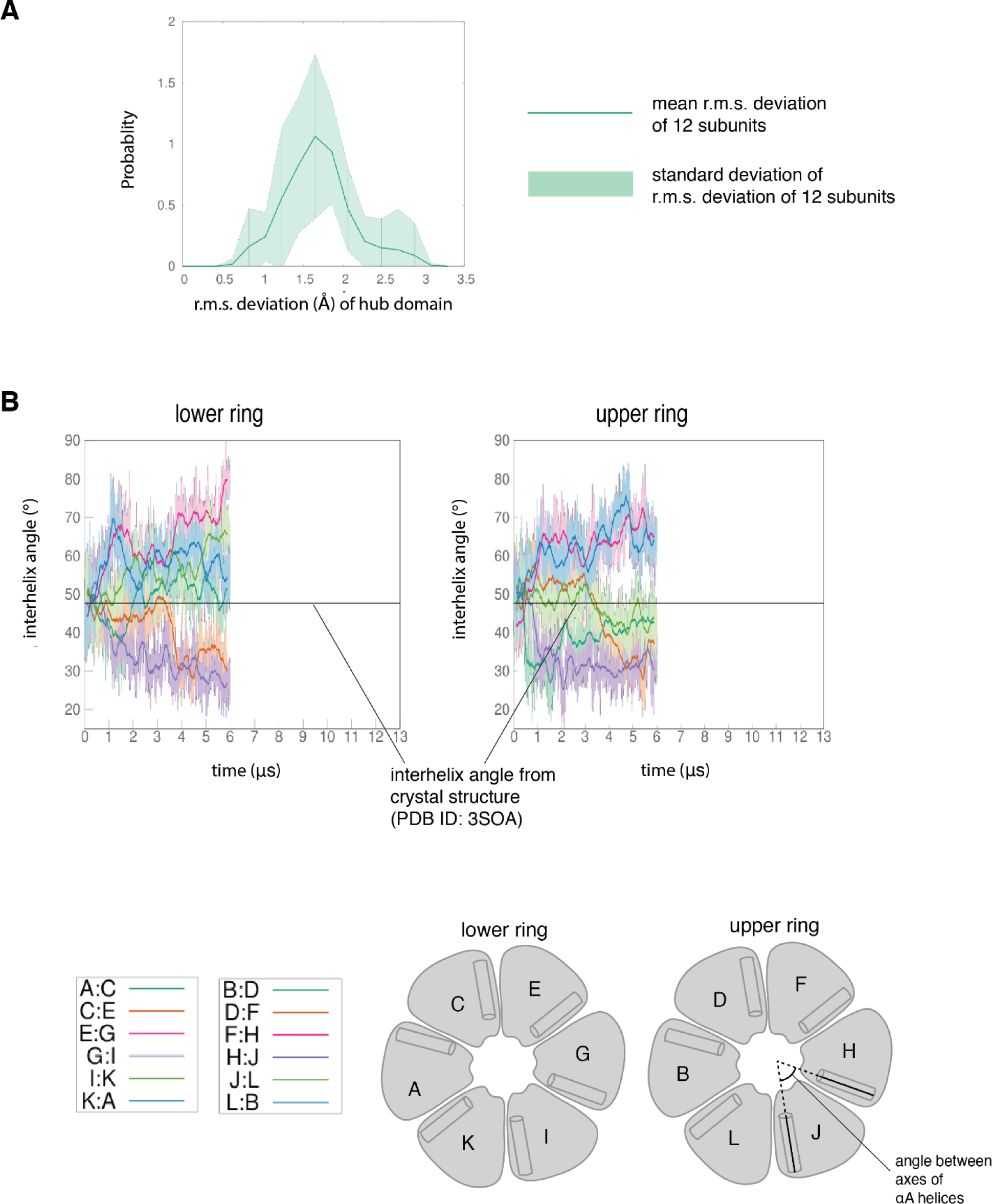
**(A)** Mean probability distribution of r. m. s. deviation of the twelve hub domains from the simulation of the dodecamer with the regulatory segments and linkers present. The pale green shaded region indicates the standard deviation of the distribution of the r.m.s. deviation of the twelve hub domains. **(B)** Variation of the angle between the axes of αA helices of adjacent subunits in the simulation without the regulatory segments. The darker traces are the time-averaged values of the interhelix angles calculated using a moving window of 240 ns while the lighter shades are the actual distances. In this simulation four interfaces in the lower ring open and the remaining two close (left), whereas in the upper ring two interfaces open and four close (right), indicating that the hub assembly is intrinsically highly dynamic. The interhelix angle between the αA helices of adjacent subunits from the crystal structure of an autoinhibited, dodecameric holoenzyme (PDB ID - 3SOA) (Chao et al, 2011), which is perfectly symmetrical, is indicated by the horizontal black line.

**Figure 8 Supplement.**
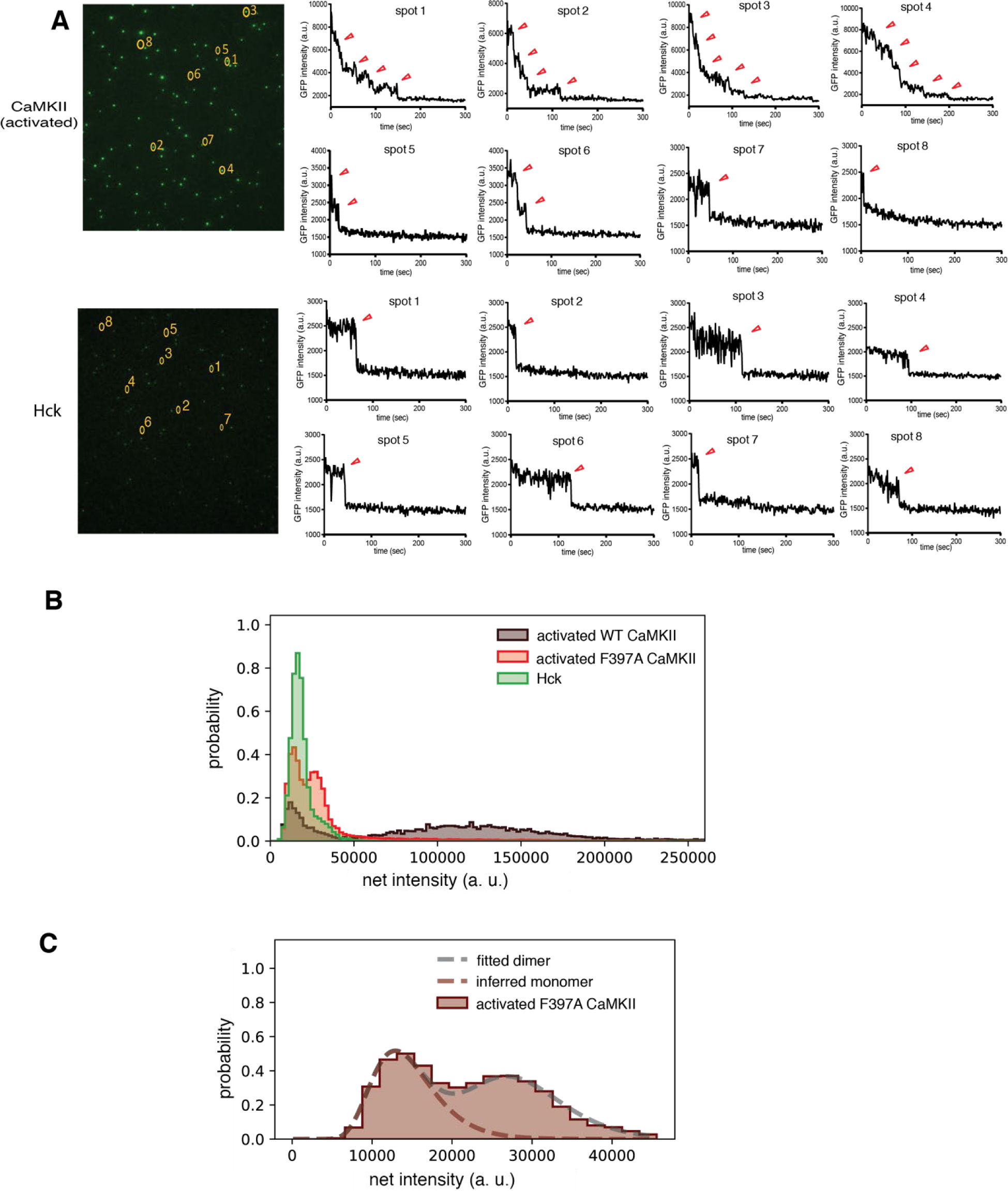
(A) Step photobleaching analyses of mEGFP-CaMKII-α (top) and Hck (bottom). One-step, two-step and multiple-step traces are observed in mEGFP-CaMKII-α. Only one-step traces are observed in Hck. The one-step traces in both samples occur at similar intensities indicating that the lower intensity spots in mEGFP-CaMKII-α correspond to monomers. **(B)** Intensity distribution curves for activated mEGFP-CaMKII-α, activated F397A mEGFP-CaMKII-α and Hck indicate that the low intensity peak observed in the CaMKII samples correspond to monomers, with the intensities that correspond to dimers appearing as a shoulder in the monomeric peak. In F397A mEGFP-CaMKII-α sample the low-intensity peak can be clearly resolved into peaks corresponding to monomers and dimers. **(C)** Intensity distribution curves generated by convolving a log-normal intensity distribution curve from the monomeric species (see Methods for a detailed description). The fit for peak at lower intensity for F397A mEGFP-CaMKII-α is shown, since this peak can clearly be resolved into monomeric and dimeric species. Here two auto-convolutions of the log-normal distribution of the monomer yields the distribution for the dimeric species. The model curve generated by twelve convolutions matches the experimental intensity distribution curve, indicating that the peak at higher intensity corresponds to a dodecamer.

**Figure 9 Supplement.**
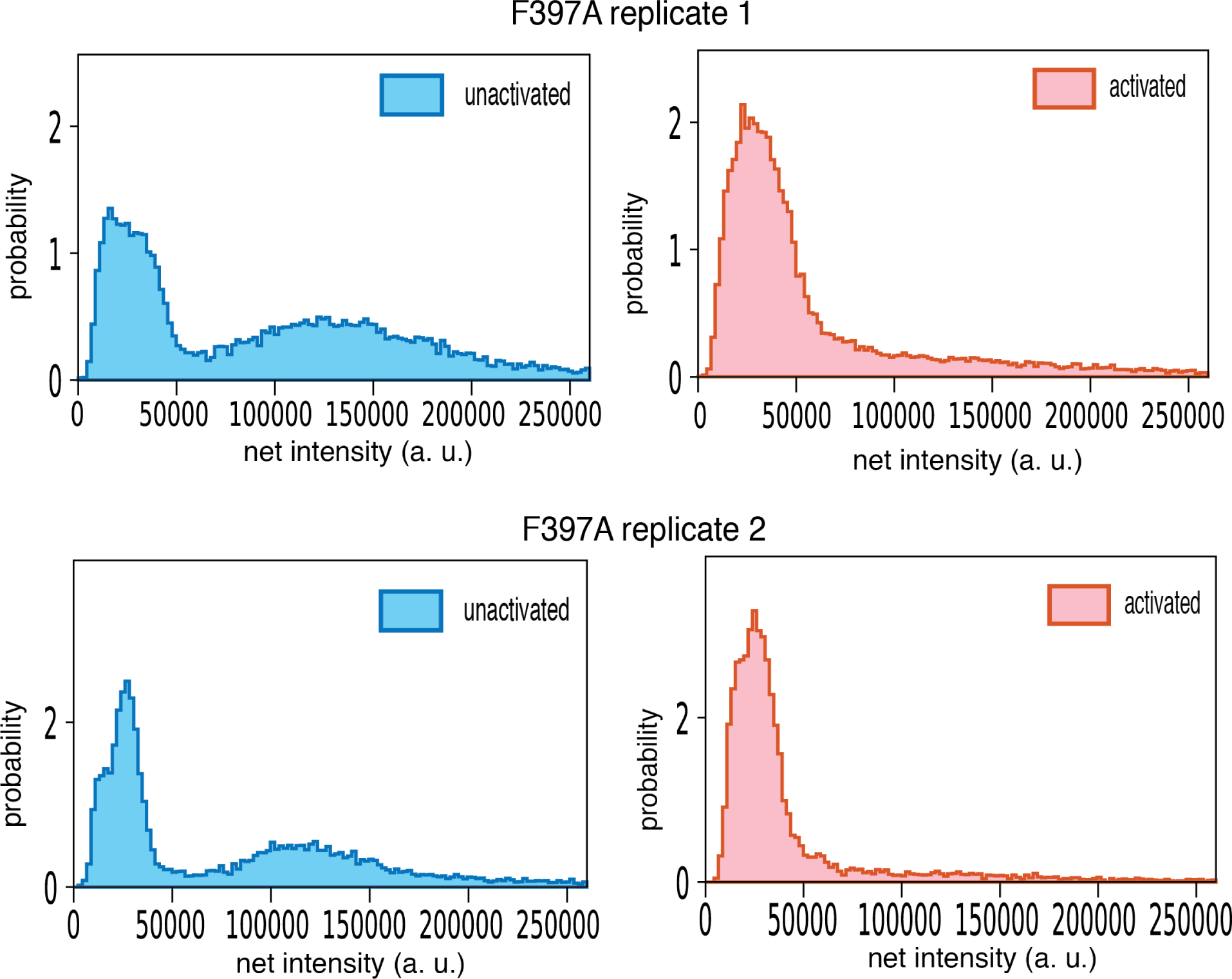
Intensity distribution of F397A mEGFP-CaMKII-α replicates. These data were collected with a 12-bit camera and the intensities were rescaled to 16-bits to be consistent with data reported in Figures 8 and 9. For these samples, a threshold value of 40 was used to exclude noisy, low-intensity spots. Only particles within a central area of 400 x 400 pixel^2^ were included in the calculation. Dark images were not collected, and the correction for uneven illumination and fringe interference effects were not done for these samples. However, both replicates qualitatively agree with the results shown in Figure 9. In both the replicates, the unactivated sample has a large peak at lower intensity that corresponds to a mixture of monomeric and dimeric species. A peak at higher intensity that corresponds to intact dodecameric holoenzymes is observed. Upon activation the area of the peaks at lower intensities in both replicates increases, while the peak at higher intensity is not observed.

